# Affective biases encoded by the central arousal systems dynamically modulate inequality aversion in human interpersonal negotiations

**DOI:** 10.1101/826529

**Authors:** Daniel AJ Murphy, Catherine J Harmer, Michael Browning, Erdem Pulcu

**Affiliations:** University of Oxford Medical School; St George’s, University of London; University of Oxford, Department of Psychiatry; Oxford Health NHS Foundation Trust, Oxford, UK

**Keywords:** Ultimatum game, computational modelling, emotional processing, social decision-making, pupillometry, neuroeconomics

## Abstract

Negotiating with others about how finite resources should be distributed is an important aspect of human social life. However, little is known about mechanisms underlying human social-interactive decision-making. Here, we report results from a novel iterative Ultimatum Game (UG) task, in which the proposer’s facial emotions and offer amounts were sampled probabilistically based on the participant’s decisions, creating a gradually evolving social-interactive decision-making environment. Our model-free results confirm the prediction that both the proposer’s facial emotions and the offer amount influence human choice behaviour. These main effects demonstrate that biases in facial emotion recognition also contribute to violations of the Rational Actor model (i.e. all offers should be accepted). Model-based analyses extend these findings, indicating that participants’ decisions are guided by an aversion to inequality in the UG. We highlight that the proposer’s facial responses to participant decisions dynamically modulate how human decision-makers perceive self–other inequality, relaxing its otherwise negative influence on decision values. In iterative games, this cognitive model underlies how offers initially rejected can gradually become more acceptable under increasing affective load, and accurately predicts 86% of participant decisions. Activity of the central arousal systems, assessed by measuring pupil size, encode a key element of this model: proposer’s affective reactions in response to participant decisions. Taken together, our results demonstrate that, under affective load, participants’ aversion to inequality is a malleable cognitive process which is modulated by the activity of the pupil-linked central arousal systems.

## Introduction

Negotiating with other people for one’s own share of finite resources is an important part of human social-economic life. Recent historic events (e.g. “trade wars” between the United States and China, Brexit negotiations between the European Union and the United Kingdom^1,2^) further highlight this importance, wherein the outcome of negotiations between only a handful of people will directly affect the lives of many others. However, computational and physiological mechanisms of such iterative social interactive decision processes remain mostly elusive.

The Ultimatum Game (UG) is a common behavioural economic measure which is used as an experimental probe of social interactive decision-making processes between two parties. In a recent study, we showed novel evidence to suggest that in iterative games *proposers* use a risky decision-making model to navigate around violating *responders’* fairness thresholds. Proposers were shown to maximise their gains by choosing between Ultimatums based on their expected returns, while taking the probability of rejection into account^3^. Rejecting an unequal proposal in the UG is also framed as a form of altruistic punishment^4-6^. Since both the proposer and the responder receive nothing when an offer is rejected in the UG, a responder’s rejection sacrifices the amount of money offered at the expense of conveying [an implicit] message to the proposer that the amount offered had been *unfair*. Consequently, the responder’s rejections in the UG violate the assumptions of the Rational Actor Model, which posits that any monetary gain is better than no gain^7^ and all offers should be accepted.

Models of inequality aversion have been particularly influential in explaining irrational choices observed in UG field experiments^8,9^. A number of subsequent neuroimaging studies^10-13^ showed neural correlates of human inequality aversion in regions associated with reward processing. On the other hand, studies using economic and evolutionary simulation models conceptualised responder behaviour in the UG in terms of reciprocity^14^, dynamic learning^15,16^, reputation building^17^ and altruistic punishment^4^. Nevertheless, reporting summary statistics of average acceptance probabilities at different offer amounts is the most common approach, and howtrial-by-trial computation of decision values underlying human responder behaviour take place in an iterative UG with an ecologically valid affective component has not been shown before.

During goal-driven social interactions, others’ facial emotions provide us a window into how they perceive our requests, helping us to decipher their otherwise hidden valuation processes. Although humans detect subtle changes in others’ facial emotions, the existing literature also suggests that people are prone to affective biases in facial emotion recognition^18-20^, which may influence their negotiation strategies. Here we will refer to these biases as “social affective bias” and will investigate the degree to which they influence human choice behaviour in an iterative UG. Although a few decision-making studies have investigated the effect of proposers’ facial emotions on human responder behaviour in the UG, it remains unknown how such affective information is integrated into decision values in iterative games. A previous study showed that responders are consistently more likely to accept offers coming from attractive faces of the opposite sex irrespective of the offer amount ^21^, while a large-scale online study suggested that offers coming from proposers with smiling faces are more likely to be accepted relative to those coming from angry faces^22^. However, a key limitation of the previous studies is the experimental approach: pairing affective faces *randomly* with different monetary offer amounts in *one-shot games*. By this methodology, on each trial, participants are asked to respond to a stimulus which is intended to be completely *independent* of what they were presented with in the preceding trial(s). Considering that in daily social interactions people’s facial emotions do not jump randomly from one affective state to another, the previous experimental approaches would only have limited ecological validity in terms of capturing real-world human social interactive decision-making processes. In order to improve on this key limitation, we designed a novel sampling algorithm for an *iterative* UG task in which proposers’ facial emotions and offer amounts were generated from two sliding windows with transition probabilities based on participant responses in the preceding trials (Figure 1). We had previously argued that this approach would break the trial-wise independence of experimental stimuli, allowing participants to experience a gradually changing social interactive decision-making environment based on their responses in the previous trials^23^. These modifications to the UG allowed us to probe participant choice behaviour using a large range of stimuli, involving both disadvantageous and advantageous/hyperfair offers.^24^ This protocol allows us to model two key influences on participant choice behaviour during social interactive decision-making: the magnitude of rewards and the proposer’s affective state. Our *a priori* hypothesis was that the proposers’ facial emotions, the offer amounts, and an interaction between these domains should influence participant’s decisions to accept or reject offers. In the following sections, using mathematical modelling of participant choice behaviour, we will describe computational mechanisms underlying how a proposer’s facial expressions influence human responder behaviour by selectively modulating perceived self—other inequality in the UG.

**Figure 1.**
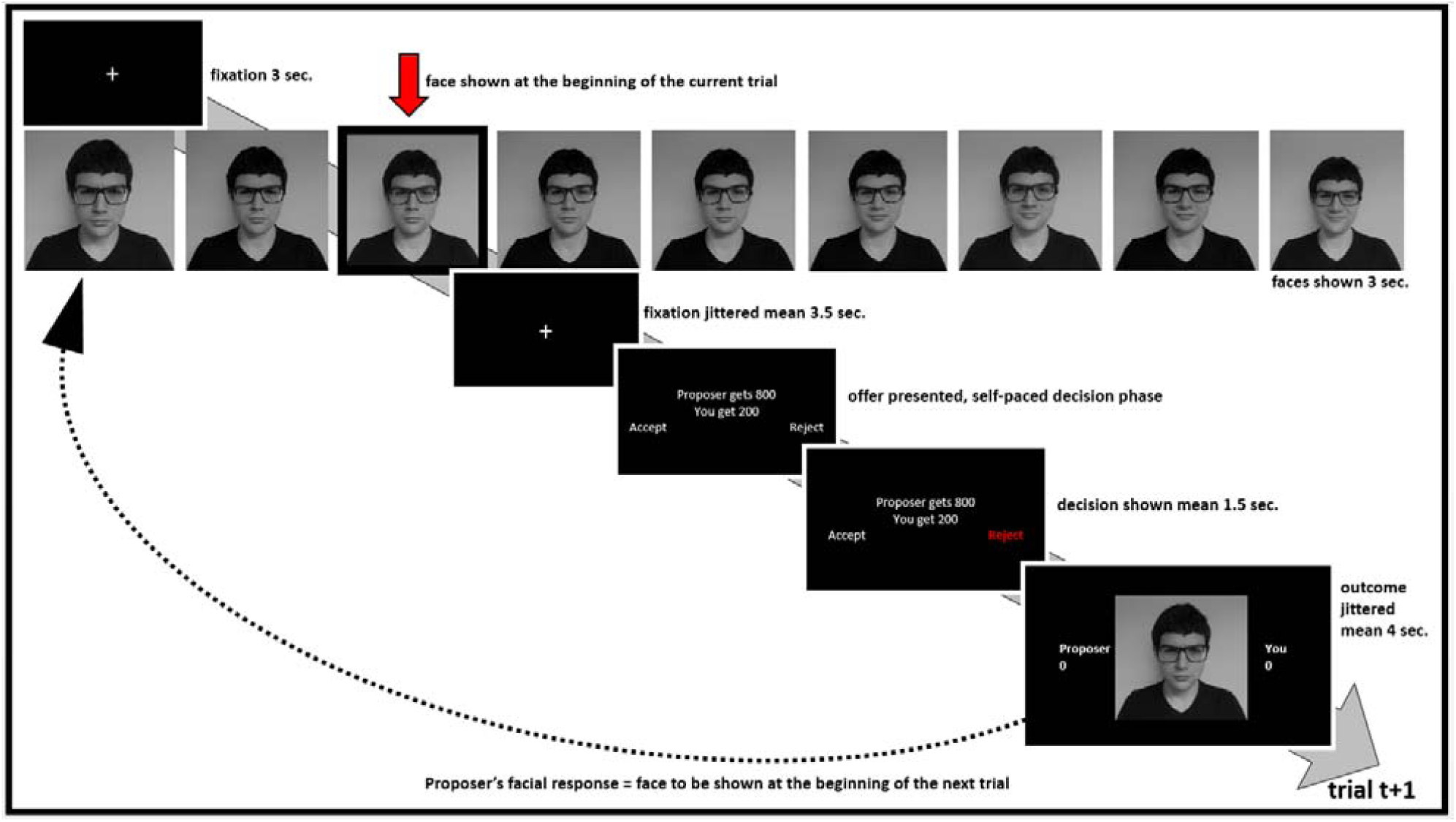
Experimental timeline of the novel Ultimatum Game task. On each trial the participant is presented with an affective face of the proposer/confederate (i.e. the red arrow) followed by an offer. In response to participant’s acceptance or rejection, the proposer’s facial emotion may stay the same or change to a neighbouring affective state based on predefined transition probabilities associated with each response type (e.g. the proposer is more likely to be happier if the offer is accepted). Similarly, based on other predefined transition probabilities, the offer on trial (t+1) may stay the same or may be revised to a neighbouring offer amount based on participant response (e.g. the offer of 200 shown in the example above for trial (t) may increase to 250 on trial (t+1), after the participant rejected the first offer). The predefined transition probabilities for facial emotions and offer amounts for the novel UG task are available in Supplementary Materials.

Changes in pupil size in the absence of any experimental manipulation of external lighting conditions is known to index the activity of the central arousal systems^25^. Previous studies highlight a role for pupil-linked central arousal systems in human reinforcement learning (RL) and value-based decision-making, particularly when performed in dynamically changing environments ^26-28^. Recent neurophysiology studies demonstrated that changes in pupil size reflect the firing rate of central norepinepheric neurons in the locus coeruleus^29,30^ (LC). These studies provide a quantitative support for a number of converging theoretical ^31^ and experimental accounts of human behaviour ^26-28,32^, all implicating a role for the central norepinephrine (NE) system in guiding behavioural adaptations in response to environmental change, which can be assessed by measuring pupil size. Nevertheless, the role of pupil-linked central arousal systems during social interactive decision-making remains unknown. Considering that our experimental task was specifically tailored to allow participants to experience an evolving interpersonal negotiation environment, we asked healthy volunteers (N=44) to perform the novel UG task while undergoing pupillometry and investigated the role of pupil-linked central arousal systems in social interactive decision-making. Informed by previous studies, we expected pupil size to encode a response to surprising offers or affective changes, as well as showing sensitivity to more subtle changes in proposers’ affective states. In line with these predictions, we present results highlighting the role of central arousal systems during social interactive decision-making, by showing that pupil size is sensitive to proposers’ affective reactions, and that it encodes a surprise response associated with offers which violate a participant’s expectations from the proposer.

## Results

### Participant Demographics

Forty-four participants recruited from the general public completed the UG experiment with pupillometry. The demographic details of this cohort are summarised in Table 1.

**Table 1.** Demographic details of participants.

| Measure | Mean (SD) |
| --- | --- |
| Age | 33.7 (11.30) |
| Gender | 68.18% Female |
| QIDS-16 | 8.11 (7.60) |
| Trait-STAI | 42.77 (13.52) |
| State-STAI | 33.43 (12.21) |

QIDS-16; Quick Inventory of Depressive Symptoms, 16 item self-report version. Trait/State-STAI; Spielberger State-Trait Anxiety Inventory. Note that scores of 6 or above on the QIDS-16 indicate the presence of mild depressive symptoms. The Trait/State-STAI has no standard cut-off scores.

### Human participants prefer fair monetary splits

Participants initially rated 100 different Ultimatum offers presented in randomised order, on a 1-9 Likert scale indicating how much they would like the proposed offer (all offers between £0.50 and £9.50 expressed in pence). These ratings allowed us to develop liking-based decision models for our main social interactive experiment. Participants’ liking ratings monotonically increased from 50p (a highly unequal offer) to 500p (a 50/50 split), where they peaked. The liking ratings were lower for offers advantageous for the participant (>500p) relative to the 50/50 split, indicating an aversion to inequality (Supplementary Figure 1).

### Nonlinearity in human facial emotion recognition

After rating the Ultimatum offers, participants rated a range of their proposer’s facial emotions, again on a 1-9 Likert scale from negative to positive. Overall, participants’ Likert ratings correlated highly significantly with the emotional valence of the proposers’ facial expressions (average correlation coefficient *r=* .869, p<.001). However, participants were significantly better at accurately rating proposer’s faces displaying positive emotions compared to negative emotions (average *r* values .821 versus .539, respectively; t(86)=-4.55, p<.001). This indicated that participants’ affective biases were more prominent for negative emotions, causing them to under-estimate the severity of negative affective displays (Figure 2A). In order to capture participants’ social affective biases, which we thought should influence the way decision values are computed in the UG, we further analysed these ratings by exploiting the properties of a two-parameter exponential-logarithmic function (see Materials and Methods, Eq.1). This novel analysis approach demonstrated that participants perceived their proposer’s affective states nonlinearly, albeit with some individual variability (Figure 2A). We further investigated whether the parameter estimates from this model were influenced by participants’ own mood states (i.e. factors such symptoms of depression and anxiety, as measured by validated clinical questionnaires). This analysis suggested that none of the continuous or categorical variables that we considered in the regression model significantly influenced estimates of the model parameters (Figure 2B). Factors such as opponent type (i.e. whether the proposer was a confederate or a computerised opponent) or opponent sex did not influence the parameter estimates either, indicating that the nonlinearity captured in participants ratings was not a direct result of the potential nonlinearity in the way proposers might have expressed their emotions while their pictures were taken initially.

**Figure 2.**
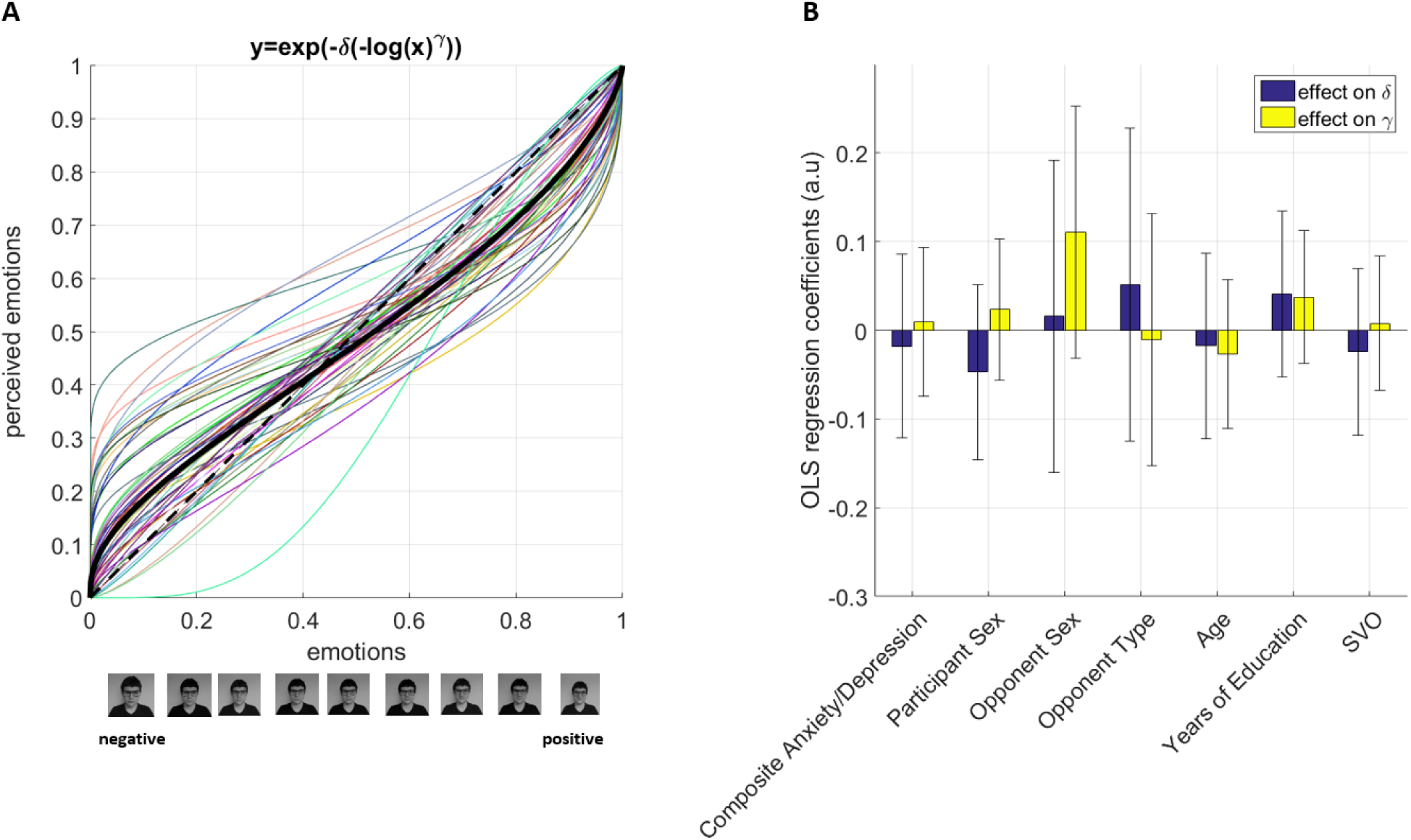
Results of the facial emotion rating task. **(A).** Participants perceived their opponent’s facial emotions nonlinearly. The x-axis shows the proposer’s faces assembled from negative to neutral to positive (as in Fig.1), and the y-axis represents participants perceived emotions according to the best fitting nonlinear weighting model. The thick black line designates the population mean. **(B)** The model parameters [δ, γ, see Eq.1] were not influenced by factors such as symptoms of depression and anxiety, or participant and proposer sex. Composite depression and anxiety scores were established for each participant by linearly transforming (i.e. summing up scores from each domain within subjects) the z-scores of depression, and state and trait anxiety measures to avoid collinearity in the model, as these measures were highly correlated with each other in this cohort of non-clinical volunteers (all r(42)>.69, all p<.001). Opponent type designates whether participants were told they would be playing against a human confederate or a computerised proposer. SVO: Social Value Orientation, which is a continuous measure defining one’s degree of prosociality. Error bars denote 95% CI.

### Proposer’s facial emotions influence choice behaviour in the UG

We first concatenated all participant data and binned average acceptance probabilities for all combinations of all proposers’ facial emotions and offer amounts (Figure 3A) and analysed this data with an ordinary least squares (OLS) regression model. This analysis suggested a main effect of facial emotion (based on t-tests on OLS coefficient estimates, t (43) =8.43), a main effect of offer amount (t (43) =8.85) and a significant facial emotion x offer amount interaction term (t (43) =3.70) influencing participants’ probability of accepting an offer (all p<.001, Figure 3B). This means that people were more likely to accept unfair offers if the proposer’s facial emotion was more positive. However, it is important to point out that these simple behavioural effects give a bird’s eye view but cannot account for the iterative nature of our social interactive decision-making task. For completeness, we report the average number of trials participants spent in each state-space during the UG experiment (facial emotion x offer amount) in Supplementary Figure 2.

**Figure 3.**
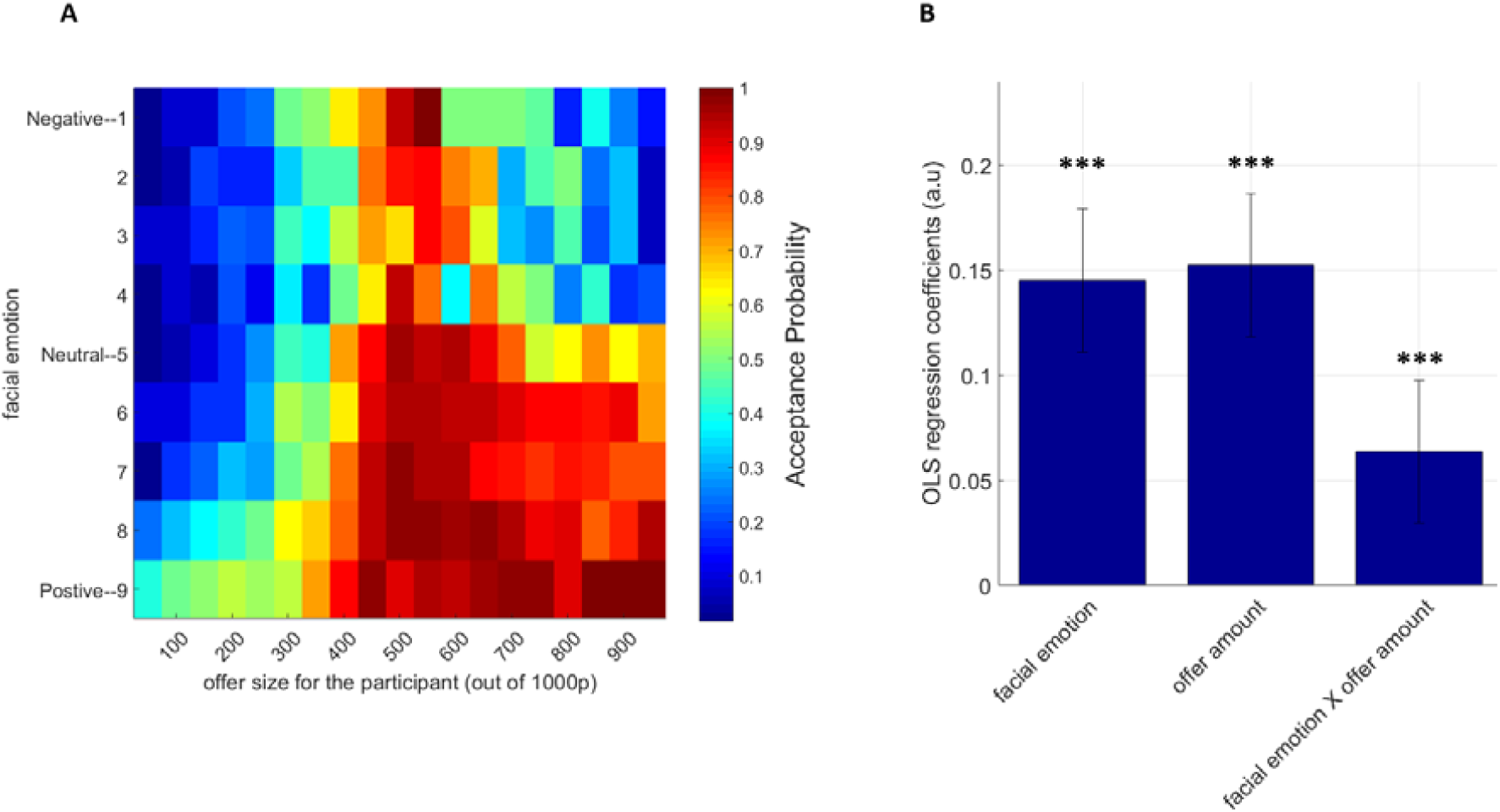
Behavioural results from the Ultimatum Game. **(A).** Participants’ average acceptance probabilities across all possible combinations of proposer’s facial emotion (y-axis) and offer amounts (x-axis) represented as a heat map. Colour bar shows the probability of accepting an offer. Changes in the colour gradient in the heat map suggests that unfair offers coming from positive faces were more likely to be accepted relative to offers associated with negative facial emotions, even if the negative-emotion offer amount is advantageous to the participant. **(B)** A formal OLS regression analysis conducted on the acceptance probabilities indicated a significant main effect of facial emotion, a main effect of offer amount and a significant facial emotion by offer amount interaction term influencing participants’ probability of accepting an offer (***p<.001).

### In iterative games recent events continue to influence participant choice behavior

One of the main features of our experimental design was that participants responded to offers generated based on their responses in preceding trials. We used this approach to break the trialwise independence of stimuli commonly employed in previous studies. Consequently, we investigated how stimuli shown in preceding trials (i.e. n-1^th^ to n-3^th^) as well as participants’ previous decisions, influenced their choice behaviour on the current trial (the n^th^ trial) using a logistic regression model. There were four regressors in this model: proposer’s facial emotion, the offer amount, a facial emotion x offer amount interaction term, and the participant’s choice. This analysis suggested all regressors from the n-1^th^ trial significantly influenced participant choice behaviour on the nth trial (all |t (43)| >2.84, p<.01), with the offer amount on the previous trial being the most significant influence (t (43) =9.56, p<.001; Figure 4). The influence of these variables on participant choice behaviour decayed further down the trials. These results are also in line with the general notion that human choice behaviour in social interactive decision-making games can be defined in terms of n-1 (i.e. memory-1) conditional-probabilistic strategies^33^.

**Figure 4.**
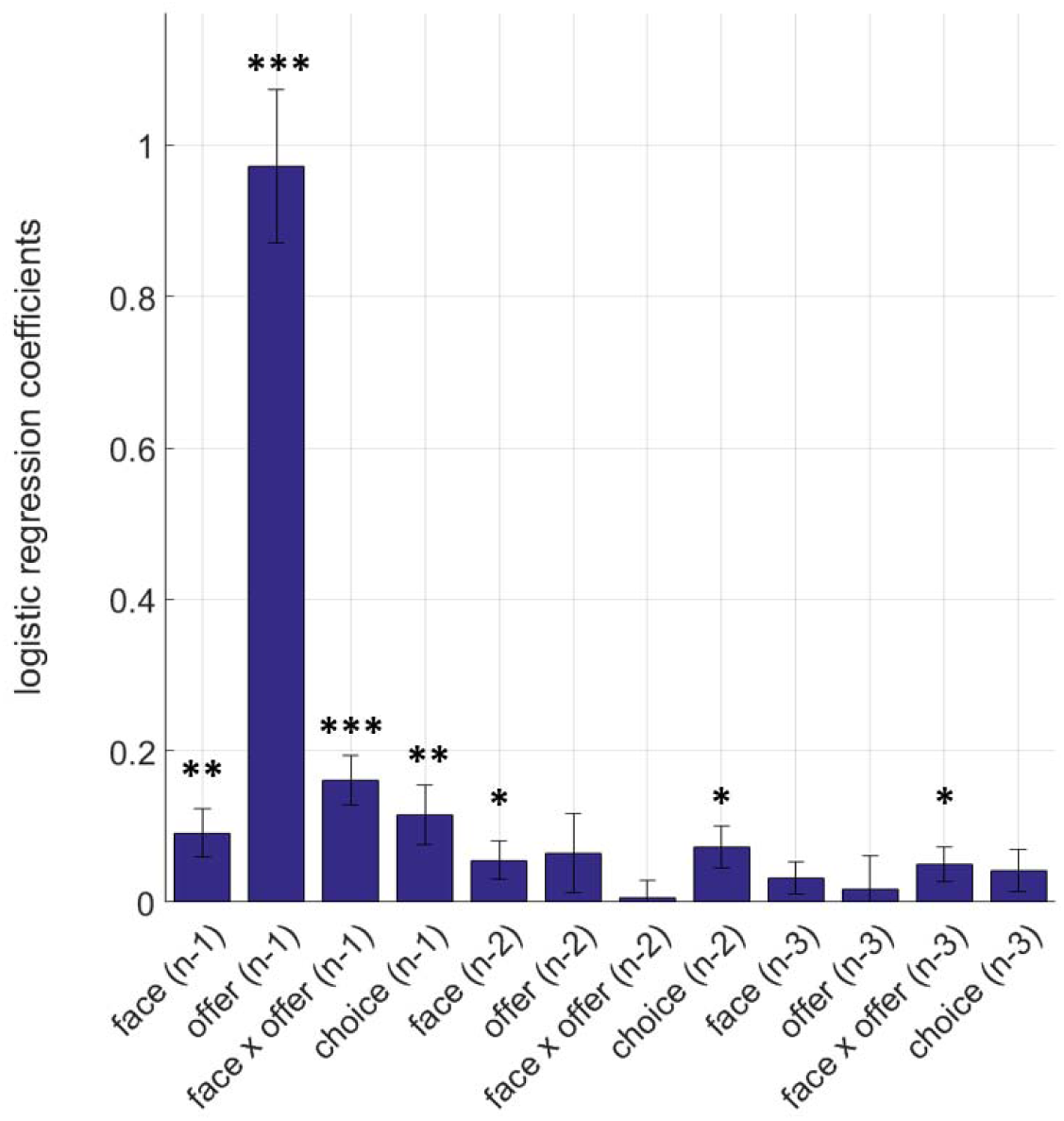
Logistic regression analysis of participant choice behaviour. Coefficient estimates from the logistic regression model fitted to participant choices on the current (n^th^) trial indicates that all regressors from the n-1^th^ trial significantly influence participant choice behaviour (***p<.001, **p<.01). Absolute values of the coefficient estimates are shown for plotting purposes. The coefficient estimates for the influence of previous choices were all negative, indicating that offers on the current trial are more likely to be accepted if offers on previous trials were rejected. The influence of these variables decayed down the trials. Error bars denote ±1 SEM.

### Proposer’s emotions dynamically modulate perceived self–other inequality

We considered a number of computational models to describe participant choice behaviour in the iterative UG task (see Methods and Materials for model descriptions). The best-fitting model (Model 7), which was able to predict participant choices in 85.8% of the trials, assumed that affective biases that modulate how the proposer’s facial emotions are perceived (as shown in Figure 2A), act on participants’ perception of inequality^9^ in the UG, and relaxes the associated negative valuation (Eqs. 1 and 9). In descriptive terms, this would mean that if the proposer insists on an offer amount which is rejected iteratively by the participant, and responds to the participant’s rejections by displaying increasingly negative emotions, at some point the proposer can overcome the participant’s negative valuation associated with perceived self–other inequality. This means that under affective load, previously rejected offers gradually become more acceptable to the responder. The best-fitting model can also flexibly adapt to other situations that occur during the game. For example, in situations where the proposer’s facial expressions are neutral, the influence of perceived affective state would is diminished (Eq. 9), whilst perceived inequality becomes relatively more important in the decision-making process. In situations in which the proposer displays increasingly positive emotions, the model then reduces the overall influence of perceived inequality on subjective decision-values, accounting for compromises (e.g. “the offer is unfair, but if I accept it, at least it makes the other side seems to be happy”). Bayesian model selection metrics^34^ supported these assertions, showing that the model in which biases of facial emotion recognition selectively act on the inequality term (Model 7)—but not on either the self-reward amounts term (Model 8) or both the self-reward and the inequality term independently (Model 9, Supplementary Figure 3)—fits better to explain participant choice behaviour. The *parabolic* (Model 7), rather than *exponential* (Model 6, Supplementary Figure 3), shape of this affective influence indicates that a proposer’s increasing negative or positive facial emotions act on the inequality term in a similar manner, while neutral facial emotions would have negligible influence on participant’s perception of inequality. Further analysis of trials mispredicted by the best-fitting model did not show any significant main effect of the proposer’s facial emotions or offer amounts (Supplementary Figure 4), indicating that the model does not fail systematically in converting these input stimuli into decision-values. However, there was a significant facial emotion x offer amount interaction in the mispredicted trials (t(43)=-3.2452, p<.01). Negative regression coefficients on this interaction term indicate that the model struggled to account for participant choice behaviour at the extremes (e.g. unfair offers coupled with very positive faces).

Similar to the OLS analysis that we reported for social affective bias parameters (Figure 2B), we analysed which variables influence the parameter estimates of the best-fitting choice model in the UG. This analysis suggested a significant relationship between participants’ social value orientation (SVO, as measured by the SVO Slider Measure^24^) and parameter estimates for the inequality term, meaning that people with higher SVO scores (i.e. people with more altruistic tendencies) perceive self–other inequality more negatively (t (43)= −2.166, p=0.036, Figure 5). Here, it is critically important to highlight that the opponent-type regressor which determines whether the participants played against a human or a computerised opponent did not have any significant main effect on human behaviour or parameter estimates. It is possible that recent advances in AI mimicking, and even excelling, human behaviour in competitive games^35-37^ might have an indirect effect on these results (i.e. human participants may find it easier to attribute human features to the computerised opponents relative to studies conducted in earlier decades).

**Figure 5.**
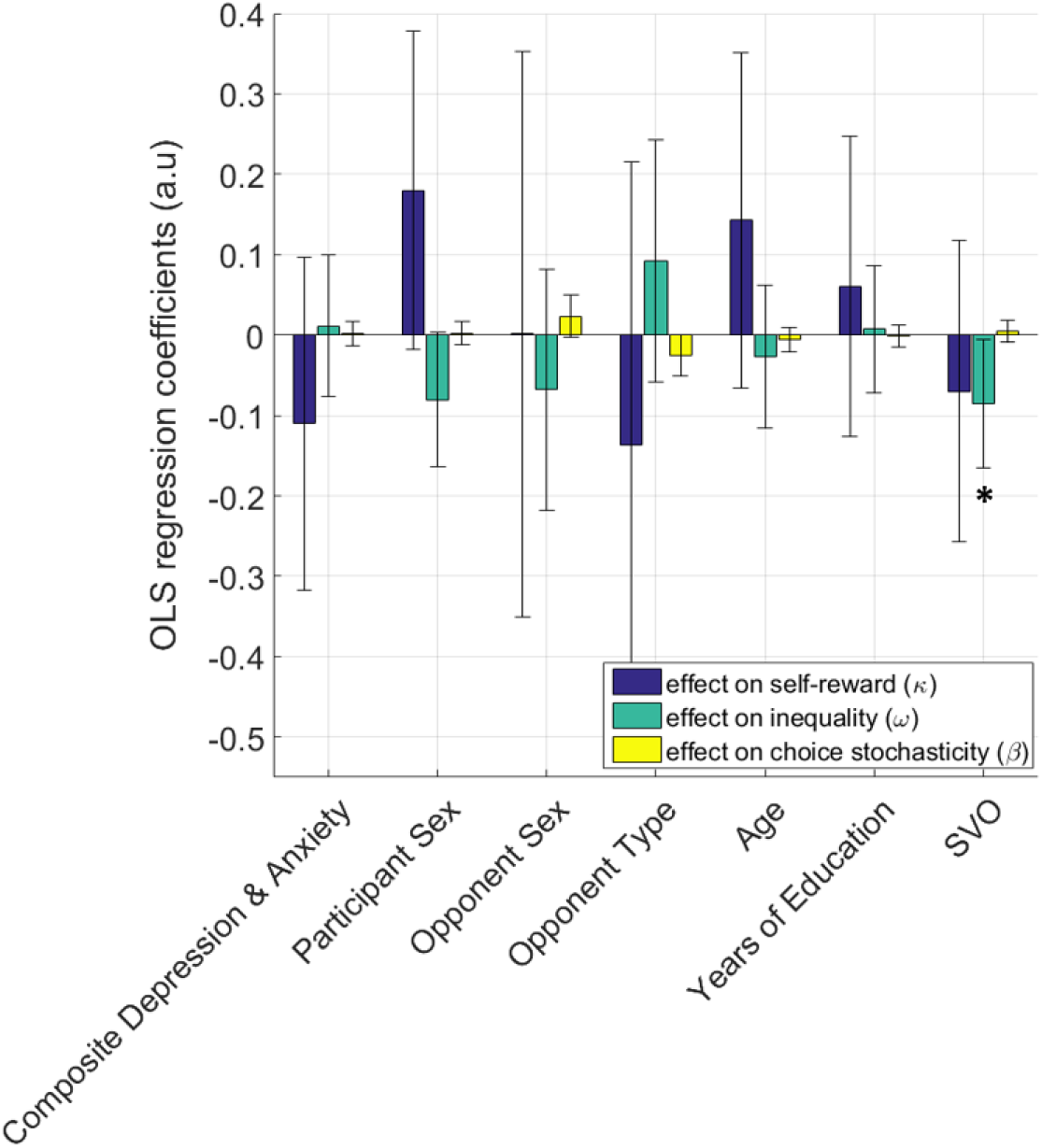
Prosocial individuals perceive self-other inequality more negatively in the UG. Coefficient estimates from an OLS regression model fitted to parameter estimates of the best-fitting computational choice model (i.e. Model 7) suggested that people with higher social value orientation will perceive self-other inequality more negatively (*p<.05). A generate-recover simulation analysis (50 iterations) based on stochastic choices generated by this model demonstrate the stability of the estimated parameters (Supplementary Figure 5). Error bars denote 95% CI. Note that these regression coefficients do not need to be subjected to further correction for multiple comparisons.

### Pupillometry Results

We analysed participants’ pupil response during the decision stage of the UG task (i.e. the window from the presentation of the offer until the end of outcome presentation). A simple model-free analysis of pupil response was performed by binning average pupil size for all combinations of proposers’ facial emotions and offer amounts (Supplementary Figure 6A), followed by an OLS regression analysis (comparable to model-free behavioural analysis reported in Figure 3B). This suggests a significant main effect of a proposer’s facial emotions on pupil size (t (42) =-2.22, p=0.028), showing that participants’ pupils dilate more when the proposer is displaying negative facial emotions (Supplementary Figure 6B).

We further analysed the pupillary data with a complementary OLS model to investigate the degree to which regressors generated under the best-fitting computational model correlated with pupil size during the UG experiment (see Materials and Methods for the details of the pupillary regression model, Eq. 11).

One possible caveat that we evaluated quantitatively before fitting the pupillary regression model is the correlation between proposers’ perceived facial emotions and the offer amounts given to the participant. Normally, one would try to decorrelate these regressors as much as possible to evaluate their unique pupillary or neural signatures. However, in the case of our task, the computerised strategy was designed to display positive faces with higher probability if the offers were accepted (as should logically happen in interpersonal exchanges in real life). In the extreme case of a proposer’s facial emotions and offers being completely decorrelated, the computerised proposer would no longer be perceived as a human proposer (e.g. proposer reacting to participant’s decisions by showing random emotions or offers). The histograms of correlation coefficients between proposers’ perceived facial emotions and decision-values generated under the best-fitting model for each participant are shown in Supplementary Figure 7A (correlation coefficient *r* (mean±SD) =.058±.26). This demonstrates that our experimental manipulation was able to deal with collinearity issues adequately, something which would otherwise compromise the quality of the pupillary multiple linear regression analysis.

In a similar manner to our previous study^3^, we also asked our participants a number of questions related to how they felt about their proposers to make sure that the computerised strategy was perceived as “human-enough” (full set of questions given in Supplementary Figure 7B legends). On average, participants were able to identify more than one person from their social circles who would make offers and display affective reactions in a similar manner to the computerised proposer (response to Q3; mean (±SD) =3.82(±.40); t (43) =7.12, p<.001), reassuring that our experimental manipulation was successful in terms of the computerised proposer adequately mimicking human behaviour while keeping the correlations between perceived faces and offer amounts within an acceptable range.

### Pupil size encodes the opponent’s affective responses which dynamically modulate perceived self–other inequality

Two key regressors that we were interested in were the decision-values generated by the best-fitting model and proposers’ facial emotions modulated by the nonlinear weighting function. Estimated coefficients for these two regressors are reported in Figure 6. Subsequent analysis (i.e. average regression coefficients binned at each second after offer presentation, analysed by one-sample t-tests from baseline) indicated that pupils dilate more in reaction to proposer’s faces which are perceived as displaying negative emotions (peak response between 1000-2000 ms, t (42)=-3.638, p=.001, Figure 6B). This result is in agreement with the model-free pupil results reported in the preceding section, and indicates that the proposer’s affective responses to participant decisions, which dynamically modulate perceptions of self–other inequality in the best-fitting computational model, is encoded by the pupil-linked central arousal systems.

**Figure 6.**
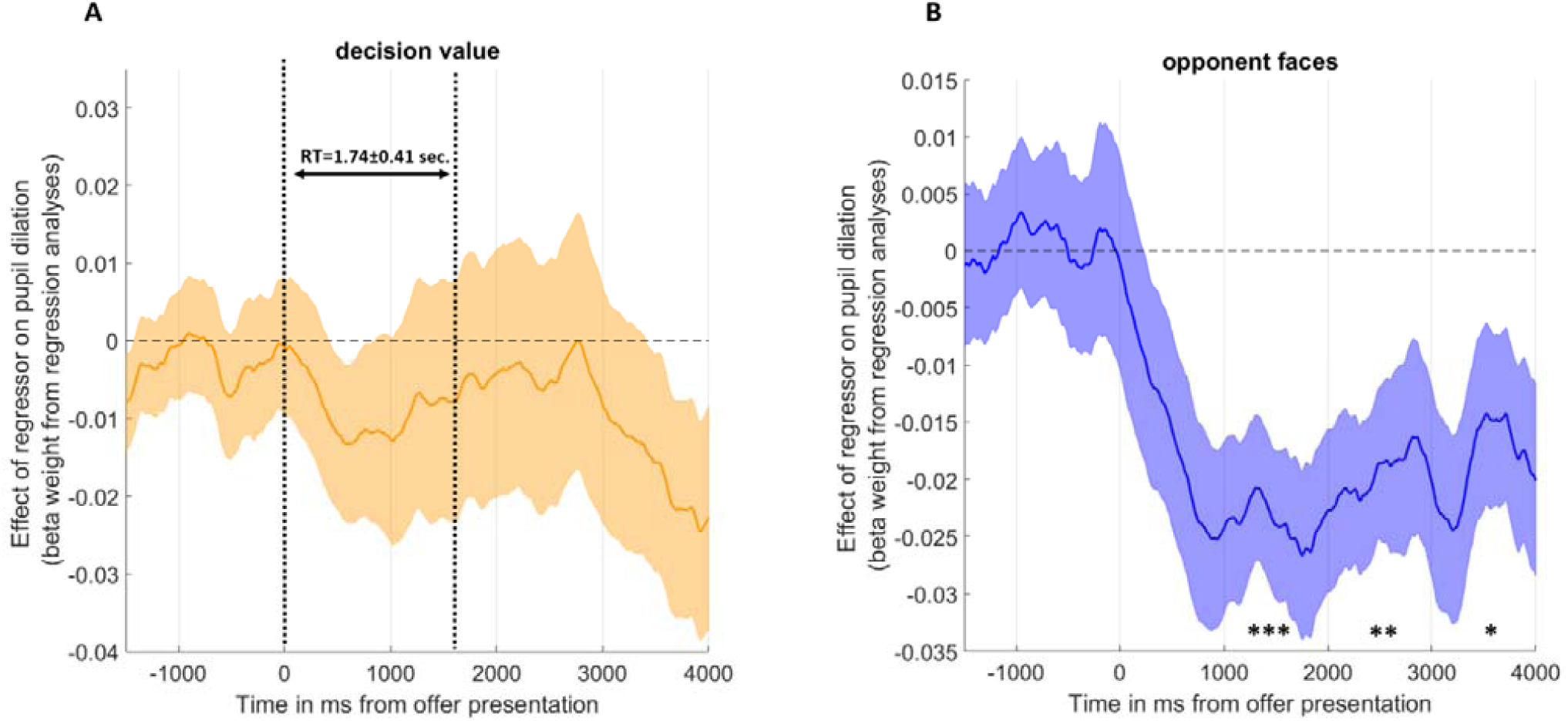
Pupillary signal of decision-values and social affective biases. **(A)** Coefficient estimates from the pupillary OLS regression model correlating with decision-values (i.e. the orange line and shading ±1SEM) from the best-fitting behavioural model. Although time evolution of regression coefficients dip during offer presentation (the window marked with dashed lines designating the decision reaction time (RT)) and post-decision interval, indicating that pupil dilate more to lower value offers, this signal was not statistically significant. **(B)** The proposer’s faces perceived as negative will lead to pupil dilation (i.e. the blue line and shading ±1SEM). The signal is statistically significant and remains significant from ∼1 second after outcome presentation (***p<.001, **p<.01,*p<.05). Error shading denotes ±1SEM.

The other regressors considered while constructing the pupil model were those quantitatively defining how the decision-making environment changed (i.e. environmental volatility and environmental noise^38^), as well as participants’ response to those changes (i.e. surprise). These were estimated from the raw stimuli (i.e. the offer amount and facial emotion valence on each trial) by a recursive Bayesian filter^38^ that can estimate the generative statistics of the stimuli uniquely for each participant, and helps with objectively quantifying how the social interaction environments change. These changes also depend on participant choice behaviour, meaning that each participant experienced a unique sequence of stimuli. The analyses of these regressors indicated that participants’ pupils dilate more in response to surprising offers (t(42) =2.166, p=.036, Figure 7A). Regressors defining environmental volatility and noise did not lead to any statistically significant pupil response, suggesting that pupil size does not encode higher order statistics of the social interactive decision-making environment.

**Figure 7.**
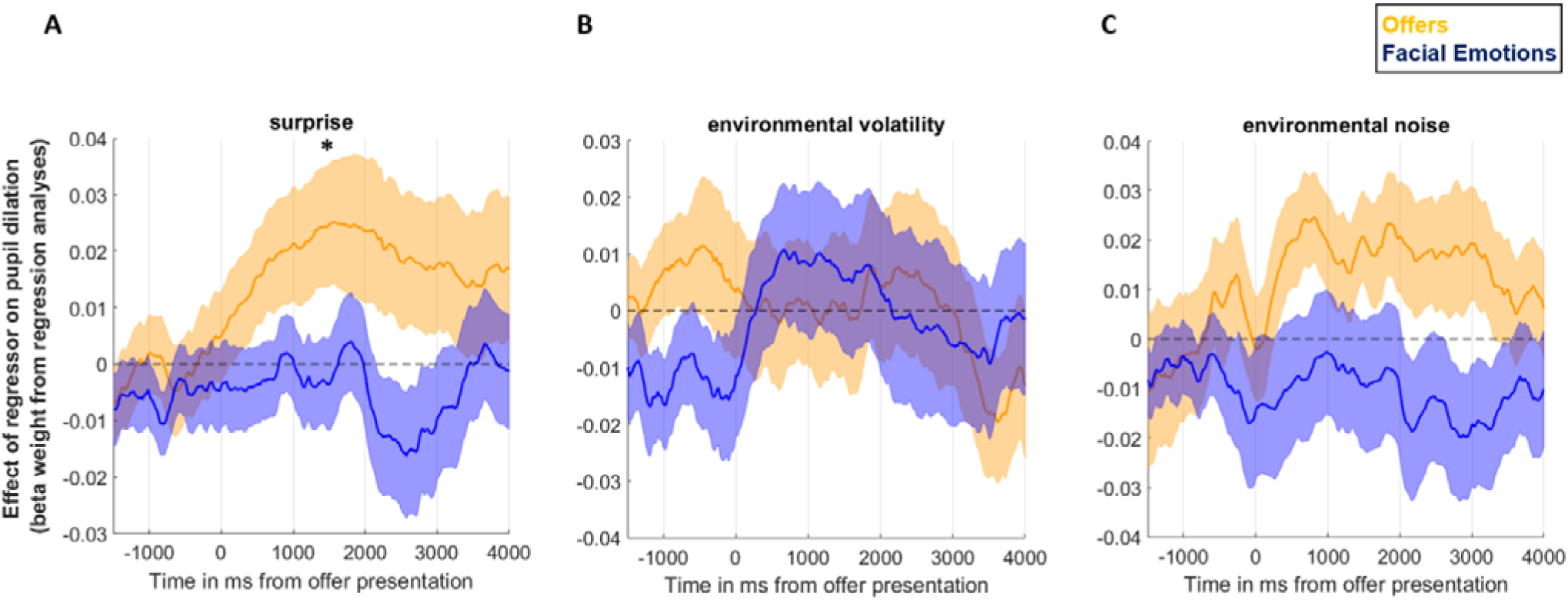
Pupillary signal correlating with regressors extracted by the Bayesian filter. **(A)** Time evolution of coefficient estimates from the pupillary OLS regression model correlating with surprise associated with offer amounts and proposer’s facial emotions. The surprise signal peaked around decision RT (*p<.05). **(B-C)** Time evolution of regression coefficients from the pupillary regression model for regressors describing environmental change. Pupil response was not statistically significant for any of these regressors. Error shading denotes ±1SEM.

### Surprising offers which violate participants’ expectations are less likely to be accepted

At the last step we investigated the degree to which regressors describing the changes in the social interactive decision-making environment which were computed by the recursive Bayesian filter also influenced participant choice behaviour. In an exploratory analysis, we fitted a logistic regression model to participant choices in which surprise, volatility and noise regressors were predictive variables (for both the proposer’s facial emotions and the offers). We further included the factors which we had an already established influence on choice behaviour (i.e. facial emotions, offer amount and their interaction as reported in Figure 3B) and the trial number (i.e. 1 to 240, total number of trials) as regressors of no interest. After excluding six extreme outliers, this analysis indicated that surprising offers were less likely to be accepted (all t(37)=-2.469, all p=.018), whereas noisy fluctuations in the proposer’s facial emotions increased participant’s probability of accepting an offer (t(37)=2.024, p=.05, Figure 8). Taken together, surprise in response to offer amounts seems to be another influence which correlates with pupil size (Figure 7A) and influences participant choice behaviour. A final control analysis investigating the correlation between the surprise regressor generated uniquely for each participant and the offer amounts observed by the participant suggested a marginally positive correlation in this cohort (correlation coefficient r(238) = .17±.09 (i.e. mean±SD), Supplementary Figure 8). This indicates that offers which violated the expectations of the participant (as estimated by a recursive Bayesian filter), but not necessarily the lower offers, were less likely to be accepted.

**Figure 8.**
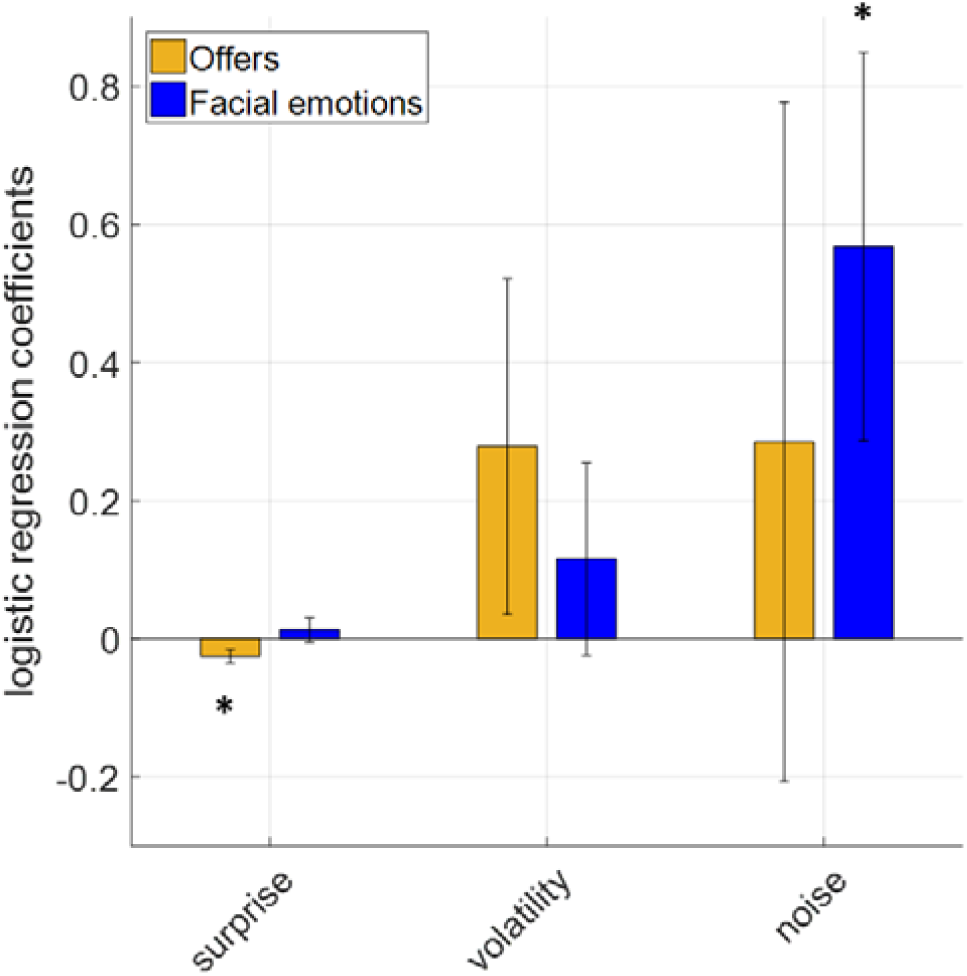
Influence of regressors extracted by the Bayesian filter on participant choice behaviour. Significantly negative correlation coefficients were observed only for the way offer amounts changed during social interactive decision-making. The results suggest that offers associated with greater surprise, environmental volatility and noise are less likely to be accepted (***p<.001, **p<.01). Error bars denote ±1SEM.

## Discussion

In the present work we describe value computations underlying human responder behaviour in an iterative UG task which also involved participants observing the proposers’ affective reactions. The results that we present demonstrate that human responders use an inequality-aversion model which is dynamically modulated by the proposer’s affective state. Our results suggest that human participants exhibit social affective biases which are represented nonlinearly (Fig. 2A). We show that proposers’ facial emotions significantly influence participant choice behaviour in social interactive decision-making (Fig. 3A-B). The influence of the proposer’s facial emotions on participant choice behaviour is also shown in results from different methods of analysing the behavioural data (i.e. both logistic regression and computational model-based). In line with previous studies, we show that parameter estimates for the inequality term from the best-fitting computational decision-making model were significantly negatively-correlated with participants’ social value orientation (SVO). This captures the intuition that people with higher prosocial tendencies perceive self–other inequality more negatively, and are less likely to accept unfair offers (Fig. 5). Computational modelling of participant choices further revealed that social affective biases dynamically modulate participants’ perception of inequality, relaxing its negative influence in a *parabolic* shape (i.e. Model 7). Participants were more likely to accept unfair offers, irrespective of whether they were advantageous or disadvantageous, when their rejections were persistently confronted with a positive or negative affective response from the proposer (also partially illustrated in the model-free faces x offer amount interaction effect, Figure 3). This model demonstrates that human inequality aversion is a malleable cognitive process, and can flexibly account for human behaviour in a number of social interactive decision-making scenarios, for example compromise behaviours to emerge in interpersonal negotiations under affective load. The pupillary regression analysis shows that proposers’ affective responses are tracked by corresponding changes in participants’ pupil size (Fig. 6B), while a surprise response estimated by a recursive Bayesian filter in response to offers given by the proposer led to pupil dilation (Fig. 7A). This indicates that pupil size also encodes violations of participants’ expectations. Subsequent analysis suggests that this surprise response is another variable which is encoded by the changes in pupil size and also exerts an influence on participant choice behaviour (Fig. 8). Thus, our data suggests that unexpected offers lead to pupil dilation and are less likely to be accepted. To the best of our knowledge, these results constitute the first line of evidence giving a detailed account of (i) how opponent’s affective responses influence value computations in social interactive decision-making; and (ii) the role of the pupil-linked central arousal systems in these value computations.

Among behavioural economic games, the UG is one of the most commonly used zero-sum game, meaning that gains are mutually exclusive between the proposer and the responder. The linear payoff structure and sequential decision-making nature of the UG makes it a good candidate for computational modelling compared to other nonzero-sum games like the Prisoner’s Dilemma (PD). The UG is known to have a strong non-monotonicity, which indicates that people’s acceptance probability does not increase monotonically with increasing self-reward magnitudes^39^. This forms the basis of non-linear aversion to inequality commonly observed in human decision-makers (Eq. 6, ω parameter). We demonstrated that our participants also behaved in this manner, as indicated by results from the analysis of choice behaviour (Figure 3A) and liking ratings (Supplementary Figure 1). Rejection of unfair offers driven by perceived inequality contradicts the Rational Actor Model, which is regarded as the optimal decision-making policy in the UG, and dictates that accepting all offers, irrespective of inequality, maximises the reward rate. Within this framework, and by using a probabilistic sampling algorithm embedded in a confederate design, we demonstrate that a modified UG task can adequately mimic social interactive decision-making in an ecologically valid manner, as supported by responses from participants on the behaviour of the computerised opponent (Supplementary Figure 7B). Our results demonstrate that the affective content of the proposers’ facial expressions serve as another source of *suboptimal* decision-making in iterative social interactions. This finding, which we demonstrate both in our model-free (Figure 3) and model-based analyses, contradicts with the predictions of the Rational Actor model, which would posit that a strategy ignoring the opponent’s affective states would be *optimal*.

The present results complement the findings of a recent work in which we described a computational model accounting for human *proposer* behaviour in the UG^3^. In these two studies, focussing on value computations underlying proposer and the responder behaviour separately allow us to restrict the vast model space and identify suitable models accounting for human social interactive decision-making. Future studies should go one step further than these preliminary reference points that we identified in the UG decision model space, and independently validate and replicate risky decision-making models accounting for proposer behaviour versus inequality aversion models accounting for responder behaviour in a two-person interactive experimental design. Although our current experimental approach tackled many of the limitations of previous UG studies, it is still far away from a naturalistic two-person exchange. Consequently, future studies involving a genuine two-person interaction would also address one of the key limitations of the current work, namely disguising an *ad hoc* probabilistic sampling algorithm within an experimental design which involves human confederates.

The extent of the decision model space has been one of the key factors limiting the number studies reporting computational models of choice behaviour in social interactive games^40-42^, as the difficulty associated with modelling recursive Theory of Mind (ToM) processes is making social interactive decision-making a topic “too hot to handle”. A recent neurophysiology study in monkeys has shown that it is possible to bypass some of these difficulties by modelling the proposer’s future decisions in a binary format (i.e. cooperate versus defect in PD) to illustrate underlying neural correlates of ToM processes^43^. However, these novel approaches require neural data with very high temporal precision, something that has not yet been implemented in human participants. As a result, and despite the fact that humans are social animals, our understanding of social interactive decision-making processes in healthy and patient groups are lagging far behind other well-defined cognitive processes such as reinforcement learning (RL)^27,28,44,45^. Nevertheless, we demonstrate that an iterative choice model converting reward and affective information into decision values can account for wide majority of iterative human social interactive decision-making (model predictive accuracy ∼86%) without recursive ToM modelling.

A number of previous studies reporting computational mechanisms which under learning and decision-making in dynamically changing environments have interpreted phasic changes in pupil size as an index for the firing of the central NE neurons^26-29^. In the current work, we showed that pupil dilation correlates significantly (i) positively with surprise associated with offers (Fig. 7A), and (ii) negatively with proposers’ affective reactions (Fig. 6B). Although previous work suggested that norepinephrine reuptake inhibitors (SNRIs) modulate neural responses to facial emotions^46^, indicating a role for the central NE system in facial emotion recognition, it is important to acknowledge that pupil dilation in response to various task components may also be under the influence of cholinergic activity in the brain (see a more detailed discussion in Faber, 2017^47^ and Muller et al., 2019^32^). Another key neurotransmitter which is involved in social decision-making^5,13,48^ and facial emotion recognition^20,49-51^ is serotonin (5-HT), and previous work has shown that pharmacological agents acting on the 5-HT system also modulate pupil dilation^52,53^. Teasing out the influence of these neurotransmitters on pupil dilation during social interactive decision-making ideally requires experimental pharmacology methods and should be tested in future studies to further understand mechanisms of causality. The current experimental approach could also be useful in teasing apart neural responses associated with information processing during social interactive decision-making, considering existing literature clearly demonstrates that facial emotion recognition and social decision-making activate fronto-striatal and limbic regions^6,10,42,48,54-56^. The main strength of the current experimental design is that it probes facial emotion processing and social decision-making domains simultaneously with a relatively fine gradient (by generating stimuli probabilistically from two sliding windows, 9 different facial emotions x 19 different offer amounts, in total 171 unique stimulus combinations, Fig. 2A), therefore it can be used as an experimental probe for addressing these mechanistic questions about information processing during social decision-making.

One of the key rating tasks which informed our modelling of participant choice behaviour in the UG evaluated participants’ social affective biases with respect to proposers’ facial emotions. We modelled participants’ ratings on a Likert scale by exploiting the properties of a two-parameter weighting function. Although our modelling approach revealed individual variability in these perceptual biases, the parameters of this model did not correlate significantly with symptoms of depression (QIDS range: 0-27) and anxiety in this non-clinical cohort (Figure 2). These results are not in line with predictions based on some of the previous studies in depressed groups which used the FERT^18,19^. Consequently, it may be worthwhile to highlight some of the experimental differences between the traditional FERT and the facial emotion rating task that we used in the current study. In the traditional FERT, participants are presented with affective faces for a very brief period, approximately for 800 milliseconds; and they are asked to label faces into one of seven different emotional categories: sad, angry, happy, surprise, fear, disgust and neutral. Brief stimulus presentation duration does not allow too much time for participants to decode the individual parts of the face while forming a judgment about its affective content. The faces are presented in an oval frame cutting out some of the facial features such as hair and ears (approximate size 56.25 cm^2^). It is possible that these aspects of the FERT create ambiguity and an information gap which probes affective biases more strongly in clinical groups relative to non-clinical volunteers. On the other hand, our rating task was self-paced (i.e. ratings were made while viewing the faces, RT (mean ± SD) = 2.90 ± 1.04 seconds). It is possible that accurately rating affective faces on a scale may be more challenging than categorising them into predetermined labels (e.g. happy, sad, fear etc.). Speculatively, ability to categorise other’s emotions quickly and accurately may have survival value similar to rapid fight-or-flight decisions. It is possible that evolution favoured agents who are better at recognising other’s facial emotions, as in the case of negative emotions, different affective states (e.g. sad vs angry) often cue distinct action tendencies. In the rating task, the faces were presented in a rectangular frame similar to a passport picture occupying a 102.7 cm^2^ area. These modifications allow participants to gather more information about the affective content of the faces, reducing ambiguity and increasing ecological validity of the stimuli and the ratings (e.g. the fact that in real life social interactions people have reasonably long time to observe others’ facial emotions, see Sonkusare, Breakspear and Guo (2019) for a critical appraisal of existing task designs in cognitive neuroscience^57^). Therefore, in future studies, it would be very important to gather large-scale data to validate the affective content of the facial expressions used in the current study, which can help understanding how symptoms of depression and anxiety influence facial emotion recognition in the wider population. While our current modelling results suggest that facial emotion processing (i.e. evaluating intensity and valence) and social interactive decision-making processes are not systematically impaired by increasing symptoms of depression and anxiety in a group of *non-clinical* volunteers, it would be informative to implement the current experimental design in clinical groups, such as patients with major depression, in which facial emotion recognition and social decision-making processes were shown to be affected in previous studies ^11,23,58,59^.

Taken together, our key results demonstrate that, under affective load, aversion to inequality in human participants is a malleable cognitive process. We show that central arousal systems—reflected in pupil size—are involved in value computations during social interactive decision-making, and track the opponent’s affective responses. We show that these perceived affective responses dynamically modulate the perceived inequality of an offer, and therefore its probability of acceptance. These findings may have important implications for understanding the cognitive processes that underlie suboptimal outcomes (e.g. compromise behaviours and settling down for an unfair split in interpersonal negotiations) in social interactive decision-making.

## Methods and Materials

### Participants

Forty-four participants were recruited from the local community via advertisements. Potential participants who had a history of neurological disorders or who were currently on a psychotropic medication were excluded from the study. In order to increase the overall generalisability of study results, eleven confederates were recruited from the staff of University of Oxford Department of Psychiatry. The confederates attended to two testing sessions. In their first visit, they were instructed to display different emotions (i.e. 9 different from negative to neutral to positive) while the pictures of their faces were taken. When their pictures were taken, the confederates were instructed to start with a neutral facial expression and then display increasing positive emotion in four increments and finally display increasingly negative emotion, also in four increments. These pictures were later used in the facial emotion rating task and the Ultimatum Game (UG, Fig. 1). In order to increase the ecological validity of these facial emotion stimuli, we did not instruct confederates to display specific emotions (e.g. disgust), but allowed confederates freedom to express negative/positive emotions as they would in their personal lives.

### Procedures

Before the experimental tasks, participants completed the Spielberger State-Trait Anxiety Inventory, Quick Inventory of Depressive Symptoms and Social Value Orientation (SVO) Slider Measure ^24^ in a pen and paper questionnaire format. After these, participants completed two rating tasks in which they were asked to rate various (i.e. in total 100 unfair, fair and advantageous offers) Ultimatum offers coming from an anonymous proposer on a 1-9 Likert scale (dislike vs liking). All offers involved splitting £10 between the proposer and the participant, and the offer amounts were presented in pence unit. After that participants rated their proposers’ (i.e. the confederate’s) pictures displaying different facial emotions, on a 1-9 Likert scale (i.e. from negative to neutral to positive, 9 different emotions in 6 iterations, a total of 54 ratings). These rating tasks were administered to establish participants’ baseline preferences independently, and the ratings were later used to construct computational models accounting for decision-making processes in the UG. The order of stimuli in both of these rating tasks were randomised for each participant in order to prevent the induction of systematic biases in perception and decision-making in the subsequent stages of the experiment.

After the rating tasks, participants completed the UG while undergoing pupillometry recording (n=44). The task consisted of 6 blocks of 40 trials each. Where available, each participant was paired with one of 11 different confederates. The participants were told that they would be playing an interpersonal negotiation game against another participant recruited from general public, and in the game they would be interacting with the other person through internet connection to an online server. To strengthen the confederate manipulation, the participants were told that their proposers attended on 2 occasions, and in their first visit they also respond to the same questionnaires as the participant did, and their pictures were taken while displaying different emotions, later to be used in the negotiation game. For some participants (47.7%) it was not possible to arrange a confederate due to feasibility issues (e.g. a mismatch between the participant, testing room and confederate availability). Those individuals were explicitly told that they would be playing against a computerised strategy which was developed based on the behaviour of a previous participant. We used this manipulation in our previous work to investigate proposer behaviour in the UG^60^. In fact, all participants played against the same computerised strategy which was developed to sample offers and facial emotions probabilistically from two independent sliding windows.

In descriptive terms, the proposer strategy was designed to test the participants’ acceptance threshold by sampling offers probabilistically around the threshold (e.g. previously rejected offers can stay the same mimicking an insisting behaviour, improve or even get lower), and displays negative facial emotions with relatively higher probability when the offers are rejected or displays positive facial emotions with relatively higher probability when the offers are accepted (details of the computer strategy is available in Supplementary Materials.). A full debriefing letter summarising the aims and objectives of the study along with reasons for deception was provided at the end of the study.

At the beginning of the UG experiment, the participants were explicitly told that their proposers would be selecting one offer out of a window of different offers to make a proposal. Participants were told that this is to make sure that their proposers could not consistently make unfair or fair (i.e. 50/50 split) offers in which case the negotiation would get stuck in a limited range of offers. This measure was taken to make sure that the decision-making process was confined to responding to combinations of faces/offers in a gradually evolving task environment, and the influence of higher order cognitive processes (e.g. Theory of Mind tracking^41^, learning about the proposer’s strategy) is limited and/or becomes redundant. We think that UG is a particularly suitable task for this purpose as it allows reducing model complexity (e.g. eliminating recursive models), relative to other tasks such as the Stag Hunt^42^, the Trust^61^ or the Inspection games^40^. This is because the Rational Actor model^7^, which posits that any gain is better than no gain and all offers should be accepted, describes the optimal strategy for maximising rewards. Secondly, recursive ToM models with which participants can try to influence the proposer behaviour would only work effectively if the proposer’s subsequent offer is more than twice as good as the offer rejected on the current trial, as otherwise the reward rate per trial cannot exceed the reward rate per trial if all offers are accepted. However, we minimised this possibility in our task design (∼9% of trials, offer amount (mean±SD): 132.6 ± 38), as offers were drawn from a sliding window and did not increase in a multiplicative manner.

Just like in any traditional UG experiment, participants were told that their task is to accept or reject these offers coming from the proposer. The participants were told that the accepted offers will be distributed as proposed, but if they reject an offer both sides would get nothing for that trial. The participants were told that after their decision, the proposer would see 9 faces displaying different emotions (i.e. the same 9 faces the participant rated previously) and would select one to communicate how s/he feels in response to the participant’s response and in the next trial the proposer would see another set of options, also giving him/her an opportunity to revise his/her offer. The participants were told that each block would start with a neutral face and a fair offer (i.e. 50/50 split) and can go any direction from that point onwards based on their negotiation ability. The participants were instructed that at the end of the game a computer algorithm would randomly select 20 trials and the outcome of those trials would be paid to each side. At this point we told participants that they may be inclined to accept all offers (i.e. the optimal decision-making according to Rational Actor model which posits that any gain is better than zero) so that those randomly selected 20 trials would always have a monetary outcome for the participant, but in that case, we told them, that their proposer may detect this tendency and try to make offers as low as possible. We then told participants to use the accept/reject responses strategically to negotiate better terms for themselves. Although our task instructions might have encouraged the emergence of choice behaviour that violates the assumptions of the Rational Actor model, it also allowed us to address our primary research questions in an unbiased manner: (i) the degree to which proposer’s affective reactions influence participant choice behaviour; (ii) how this affective information is incorporated in decision values in iterative games.

During pupillometry recording, participants’ heads were stabilised using a head-and-chin rest placed 70 cm from the screen on which an eye tracking system was mounted (Eyelink 1000 Plus; SR Research). The eye tracking device was configured to record the coordinates of both of the eyes and pupil area at a rate of 500 Hz. The pupillometry data collection lasted approximately 70 min per participant.

### Modelling affective biases in facial emotion recognition

We modelled participant rating responses to affective faces (on a 1-9 Likert scale from negative to positive) by exploiting the properties of the 2-parameter probability weighting function^62^:

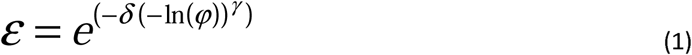

where *φ* is the true emotional state of the proposer’s face as displayed, and can take values from 0.1 to 0.9. The parameters [*δ, γ*] determine the curvature of the weighting function and where it crossed the *ε* = *φ* diagonal line (Figure 2A). We implemented the probability weighting function like a nonlinear regression model that minimises the difference between model estimates and participant’s ratings (after performing a simple linear transformation by dividing the Likert ratings by 10, such that both the ratings and the estimates are bounded by 0 and 1).

### Modelling liking ratings for Ultimatum Offers

In line with previous literature^54^, we modelled participants liking ratings (*χ*) for each Ultimatum Offer (again, based on participant ratings on a 1-9 Likert scale) with a multiple linear regression model.

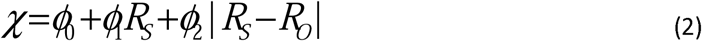

where R_S_ is the self-reward and R_O_ is the reward amount to the proposer, and absolute value difference accounts for how much the participant cares for inequality. Here, it is important to clarify that because all offers involve splitting £10, R_O_ is not entered to this regression model as an independent regressor as all monetary information is conveyed by the self-reward amount and the inequality term.

### Modelling participant choice behaviour in the Ultimatum Game

The model-free analyses of participant choice behaviour indicated that both offer amount and the proposer’s facial emotions should influence how people generate decision-values in the UG. Here, we formally describe all the models that we considered, to identify the cognitive model which accounts for participant choice behaviour the best.

The first model that we considered assumes that participants construct the decision-value 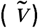 of each offer utilising a Constant Elasticity of Substitution (CES) function, commonly used to account for consumer behaviour ^63,64^. According to the CES function, the decision-value is computed as:

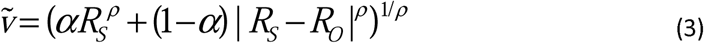

where *ρ* is a nonlinear power utility parameter determining the concavity of preferences and *α* determines how much weight is assigned for the self-reward magnitude or the absolute value difference between the self and the other’s reward magnitude (i.e. the inequality term).

We considered another model in which the decision-value is generated by comparing the inequality term relative to one’s fairness threshold:

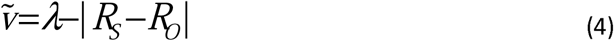

where *λ* is the fairness threshold parameter freely estimated between 50 and 950 (i.e. bounded by the minimum and maximum offer amount), defining the participant’s subjective threshold in term of what they regard as acceptable.

The next model assumed that participants accept or reject offers based on their liking ratings, as established by the independent rating task participants completed prior to the social interactive decision-making task. This model allowed us to validate previous accounts of value-based decision-making which demonstrated that choice preference does not always align with participant ratings^65^.In this model, participant’s liking of offers during the UG is estimated by feeding each participant’s estimated coefficients from the linear regression model back to Eq.2. Here, the decision-value is equal to:

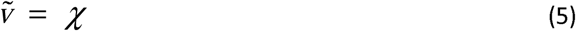

We also considered another model with a similar structure in which the decision-values are generated online during the social decision-making task, instead of depending on participant’s previous liking ratings. This model addresses the prediction that there will be a dissociation between how much people like offers when these ratings do not have any financial consequence and how they value them in a social interactive context with monetary consequences (i.e. the fact that participants would be paid the outcome of 20 randomly selected trials). Here, the decision-value is computed as

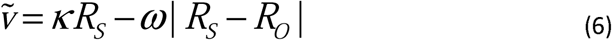

For model simplicity we used a single inequality aversion parameter, instead modelling advantageous and disadvantageous inequality separately. All models assume that conditions with relatively higher subjective value should be more likely to be accepted and participants’ acceptance probability is generated by a sigmoid function:

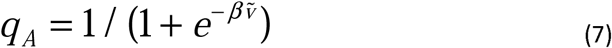

where *β* is the inverse temperature term which modulates the stochasticity of participant choices.

We first chose between these models which define participants’ likelihood of accepting an offer solely based on the numerical components of the offer amount. We implemented this reduction in model space based on our model-free analysis which suggested that offer amount has a greater influence on acceptance probabilities than the proposer’s facial emotions. Group-wise sum of Bayesian Information Criterion (BIC) scores indicated that the model described by Eqs. 6 and 7 was the best fitting [inequality aversion] model to account for how offer amounts were translated to decision-values.

We then used this best fitting model from the first stage as a template to further evaluate the degree to which facial emotions of the proposers dynamically influence the way decision-values are computed in the social decision-making task on a trial-by-trial basis. Based-on previous literature, we considered a model in which the effect of proposer’s facial emotion depends on the participants’ individual variability in how malleable they are to external emotional influence ^66^:

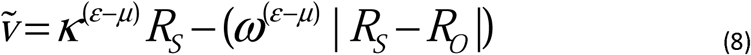

where parameters κ and ω are estimated between 0 and 10 and determine the degree to which the modulations of self-reward and the inequality term are subject to emotional influence. In this model, social affective biases (Eq.1) act like an exponential function to modulate this influence parameter [6] and μ is the value where perceptual biases crossover the diagonal line (Figure 2A, μ=0.4). We considered 3 variants of this model where proposer’s facial emotions influence the self-reward amount (R_S_) or the inequality term (|R_S_ – R_O_ |), or both independently.

We also investigated whether the proposer’s facial emotions influence the inequality term in a parabolic, rather than an exponential functional form:

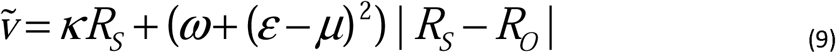

For any given offer amount, this formulation relaxes the negative modulation of inequality as the proposer’s facial emotions are gradually getting negative or positive, while assuming that neutral faces would have limited influence on the inequality term.

Finally, we considered a competing model which assumes that the proposer’s facial emotions influence the value of offers through a weighted integration.

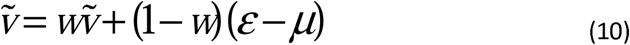

Here, *w* is a free parameter estimated between 0 and 1, and determines the degree to which the participant assigns credit to subjective value of the offers and/or to the proposer’s facial emotions which are nonlinearly modulated by affective biases. The additive integration means that other’s facial emotions should have an intrinsic value, an assumption that is in line with existing literature ^67^.

All model parameter estimation followed our existing protocols^26,28,45^, namely by a Bayesian model fitting procedure, by calculating the full joint posterior distribution of the parameters over the whole parameter space, and deriving exact parameter values by integrating these probabilities with their corresponding discrete parameter values.

### Pupillometry data preprocessing

Eye blinks were identified using the built-in filter of the Eyelink system and were then removed from the data. A linear interpolation was implemented for all missing data points (including blinks). The resulting trace was subjected to a low pass Butterworth filter (cut-off of 3.75 Hz) and then z transformed across the session (Browning et al., 2015; Nassar et al., 2012). The pupil response was extracted from each trial, using a time window based on the presentation of the offer amount. This included a 7.5 second baseline period before the presentation of the outcome (including the period at the beginning of a trial marked by a fixation cross and presentation of the proposer faces alone, Figure 1), and a 4.5 second period following offer presentation. Baseline correction was performed by subtracting the mean pupil size during the 7.5 seconds baseline period prior to the presentation of each offer, from each time point in the decision and outcome period^68^. This baseline correction, which also included the first facial expression shown within a trial (see Figure 1), allowed us to extract the phasic pupillary response and controls for potential fluctuations in luminosity within each trial. Individual trials were excluded from the pupillometry analysis if more than 50% of the data from the outcome period had been interpolated (mean =10.9% of trials) (Browning et al., 2015). The preprocessing resulted in a single sets of pupil time-series per participant containing pupil dilation data for each of the included trials.

### Regression analysis of pupillometry data

We implemented an OLS model in a similar manner to a functional neuroimaging (fMRI) analysis, by fitting the regression model to the pre-processed pupillary data at every 2-millisecond time point. The pupil model had 10 regressors as defined by the following linear regression equation:

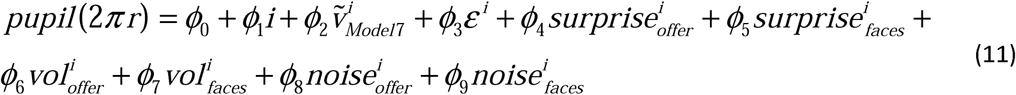

Here we provide brief descriptions for each regressor in the model. The regressor for the constant term (ϕ_0_) captures the variance in pupil size across all trials in the task. The regressor (ϕ_1_) accounts for the effect of trial numbers, as a proxy for accumulated fatigue (*i*: taking values from 1 to 240). We entered one regressor (ϕ_2_) which represents the subjective value of each offer under the best-fitting behavioural model. The regressor (ϕ_3_) encodes the perceived facial emotions modulated by 2 weighting parameters (Eq.1). The rest of the regressors were defined by a recursive Bayesian filter that we recently reported^27^ which can optimally track the hidden structures (i.e. environmental volatility,vol in the above formula, and noisy fluctuations, noise, in the environment) of dynamically changing environments. The surprise regressor was congruent to the conditional –log probability of an offer coming from a distribution with mean and standard deviation (SD) estimated by the Bayesian filter. This would mean that stimuli which violate the expectations of the observer should lead to a greater pupillary surprise signal. This regressor was calculated for both the offers and the proposer’s facial emotions. All regressors were demeaned prior to model fitting. Fitting a multiple linear regression model to pupillary data is akin to analysis of fMRI datasets and allows regressors to compete for variance in the data. Therefore, resulting regression coefficients no longer need to be subjected to further correction for multiple comparisons by the number of regressors in the model.

## ACKNOWLEDGEMENTS

This study was funded by National Institutes of Health Research (NIHR) Oxford Biomedical Research Centre funding allocated to EP. DAJM and EP collected the data. EP analysed the data. MB developd the recursive Bayesian filter. All authors contributed to writing up of the manuscript. The authors would like to thank Dr. Alexander Kaltenboeck for his help with the study. The views expressed in the manuscript are those of the authors and not necessarily those of the NHS, the NIHR or the Department of Health. MB is supported by a Clinician Scientist Fellowship from the MRC (MR/N008103/1) and by the NIHR Oxford Health Biomedical Research Centre. He has received travel expenses from Lundbeck for attending conferences and acted as a consultant for Jansen Research and CHDR. CJH has received consultancy fees from p1vital. Lundbeck, Pfizer, Sage Pharmaceuticals, Servier, Zogenixs and J&J.

## Supplementary Figures

**Supplementary Figure 1.**
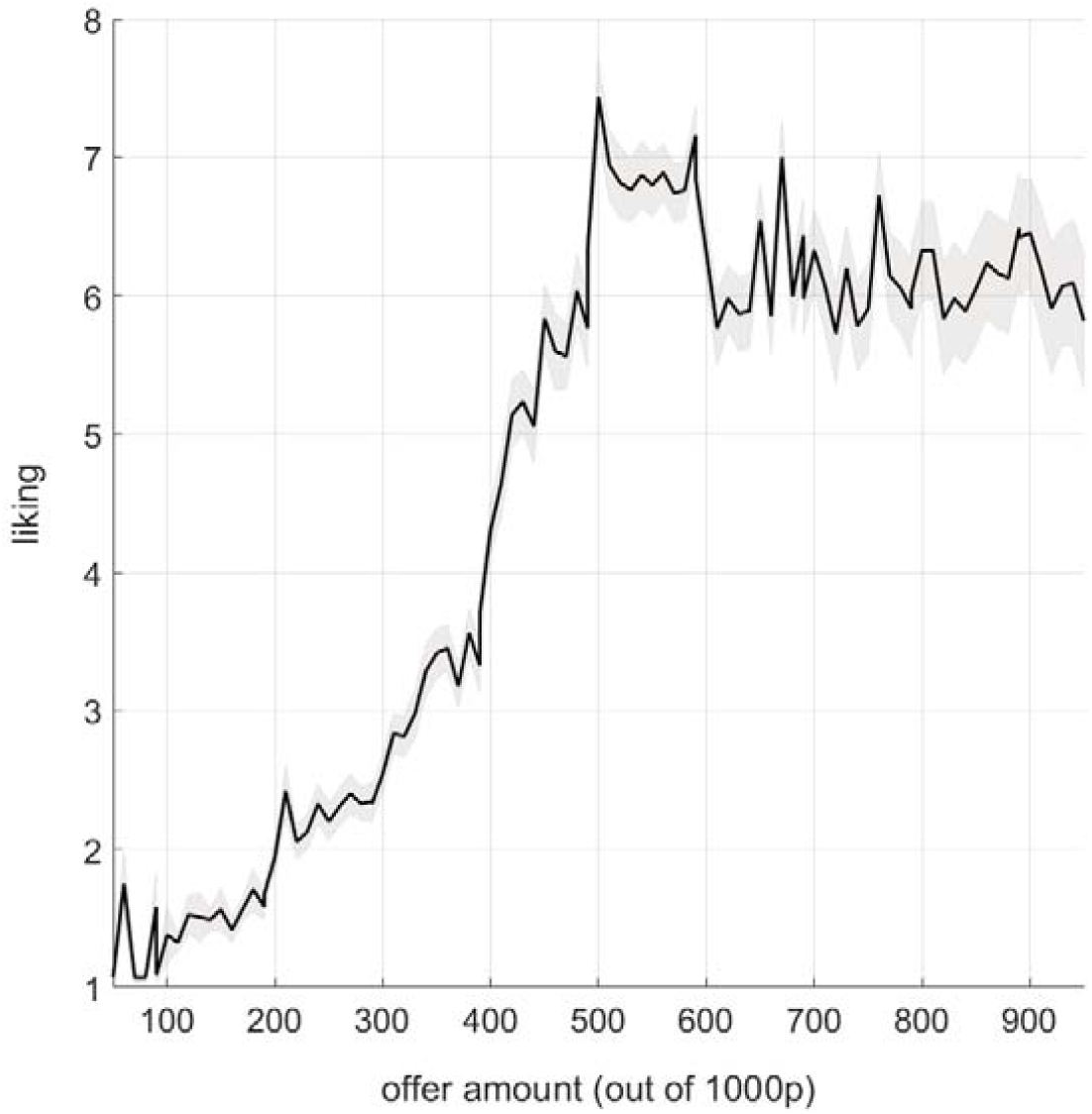
Participants’ liking ratings monotonically increase from 50p to 500p, which was the point of 50/50 split in the current experimental design. Relatively lower ratings for advantageous offers (>500p) indicate inequality aversion in this cohort. Wider SEM shading around the ratings of advantageous offers indicate a greater individual variability in terms of self-other inequality relative to how unfair offers were rated.

**Supplementary Figure 2.**
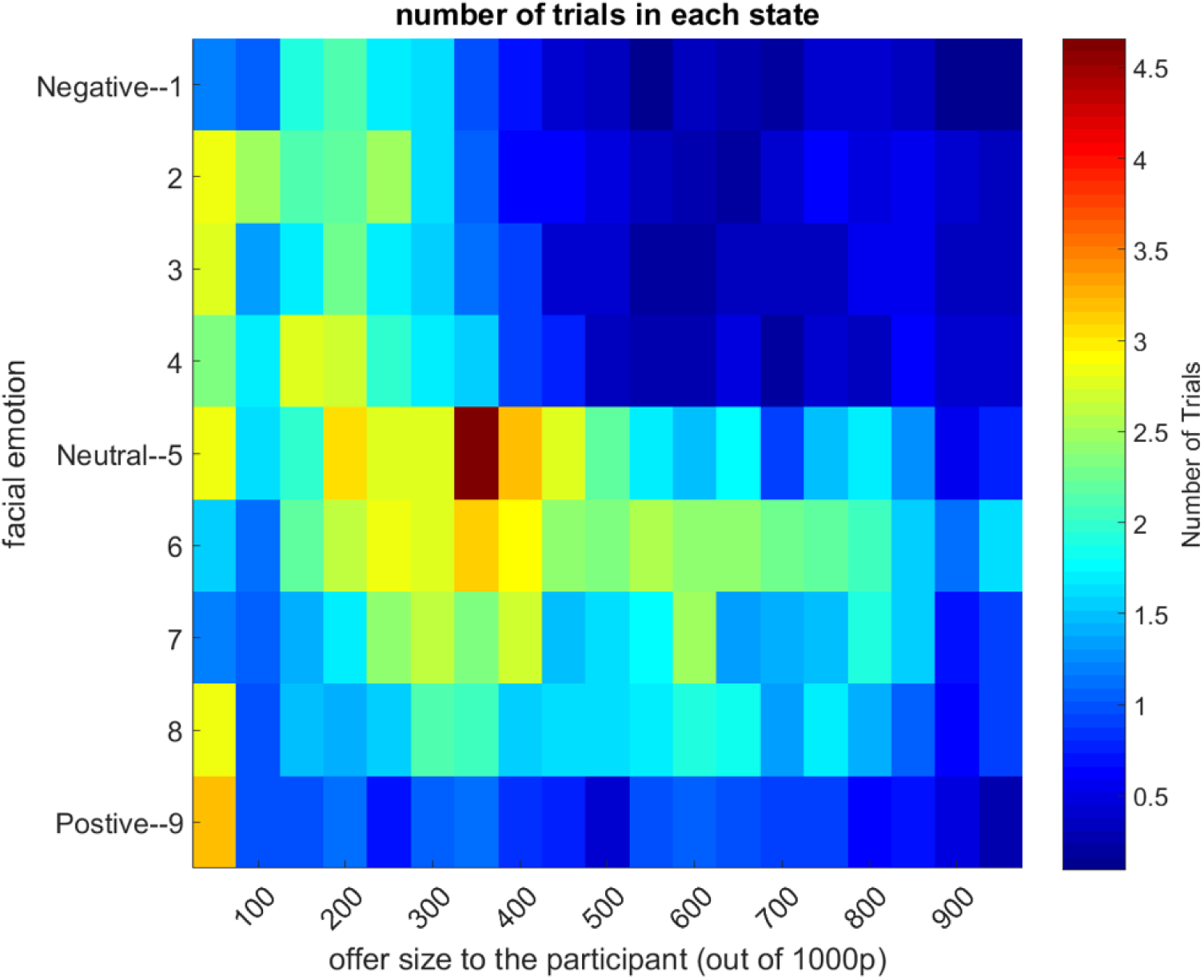
Out of 240 trials, average number of trials participants spent in each state-space (i.e. combinations of proposer’s facial emotion (y-axis) and offer amounts (x-axis)) represented as a heat map. Colour bar shows the number of trials. On average participants had 12.6±15.9 (mean±SD) trials in the upper right hand quarter (i.e. advantageous offers while the proposer is displaying a negative facial expression).

**Supplementary Figure 3.**
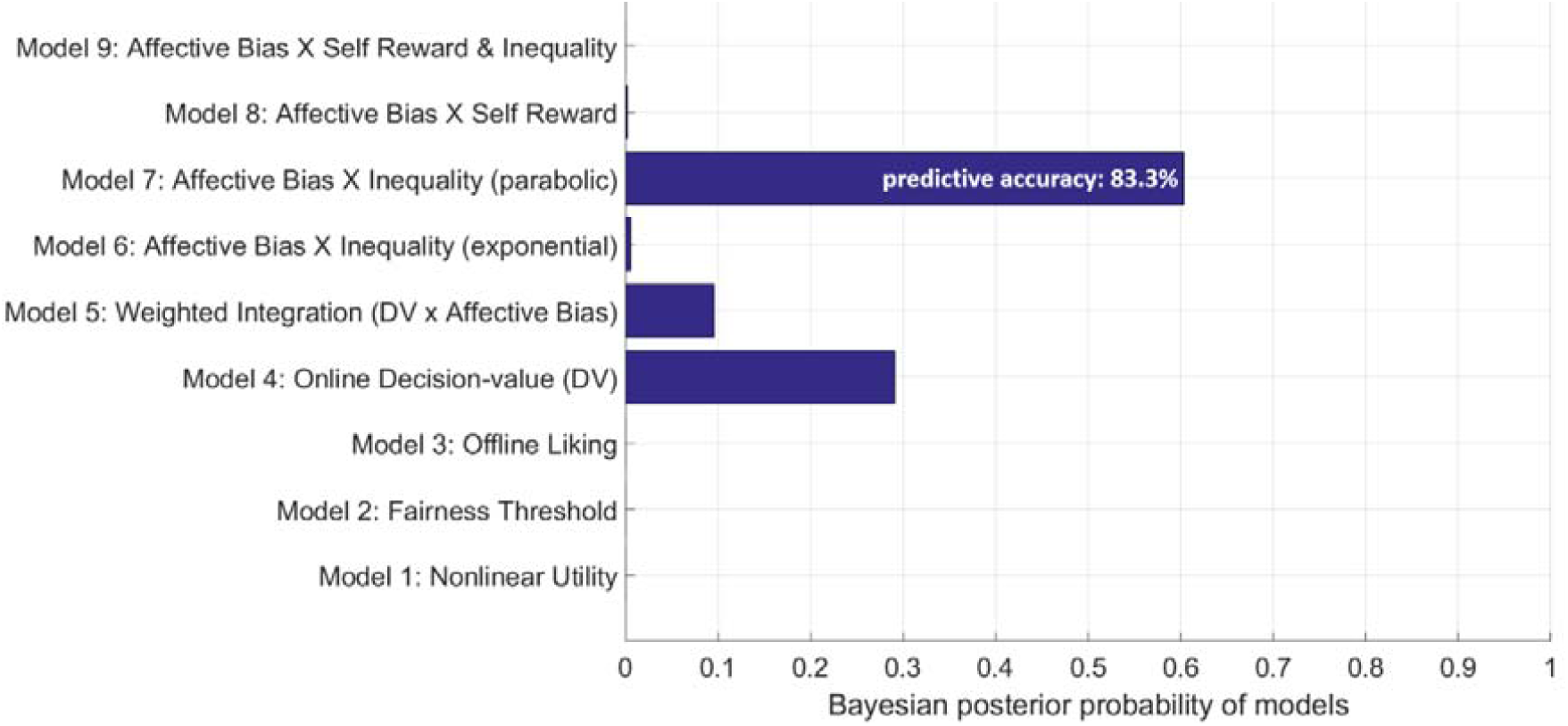
Model selection implemented by feeding each model’s log-likelihood values in a matrix format (number of participants x number of models) to spm_BMS.m (i.e. the Bayesian Model Selection from SPM12 library), suggests that Model 7 is most likely to be the generative model for the observed data (based on model exceedance probabilities). This model suggests that social affective biases selectively act on the inequality term in a parabolic form.

**Supplementary Figure 4.**
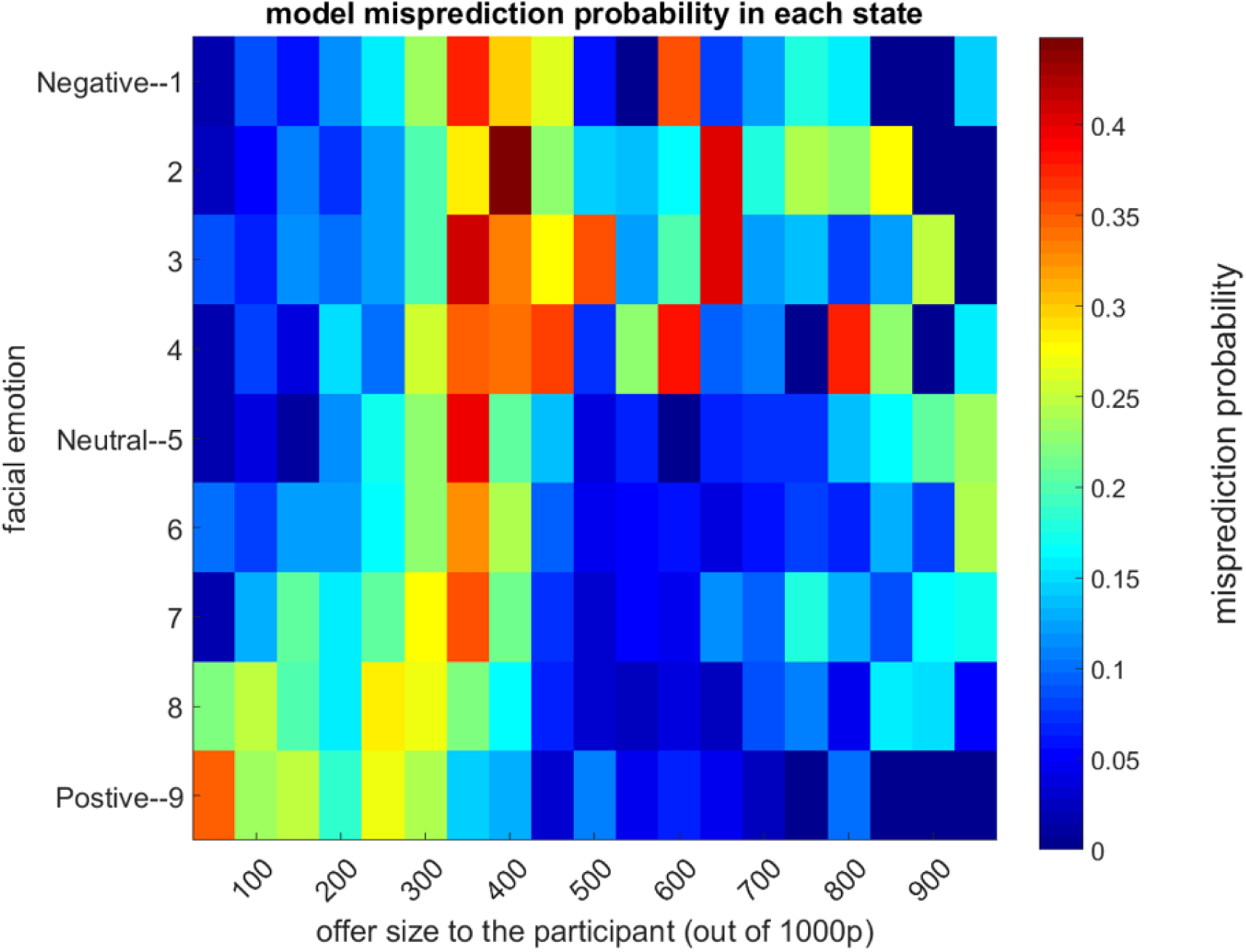
Misprediction probabilities of the best-fitting model over the stimuli space. A regression analysis of the trials mispredicted by the best-fitting model did not show a significant main effect of proposer’s facial emotions or the offer amount, indicating that the model does not miss the behavioural effect of faces and the offer amount (Figure 3) in a systematic manner.

**Supplementary Figure 5.**
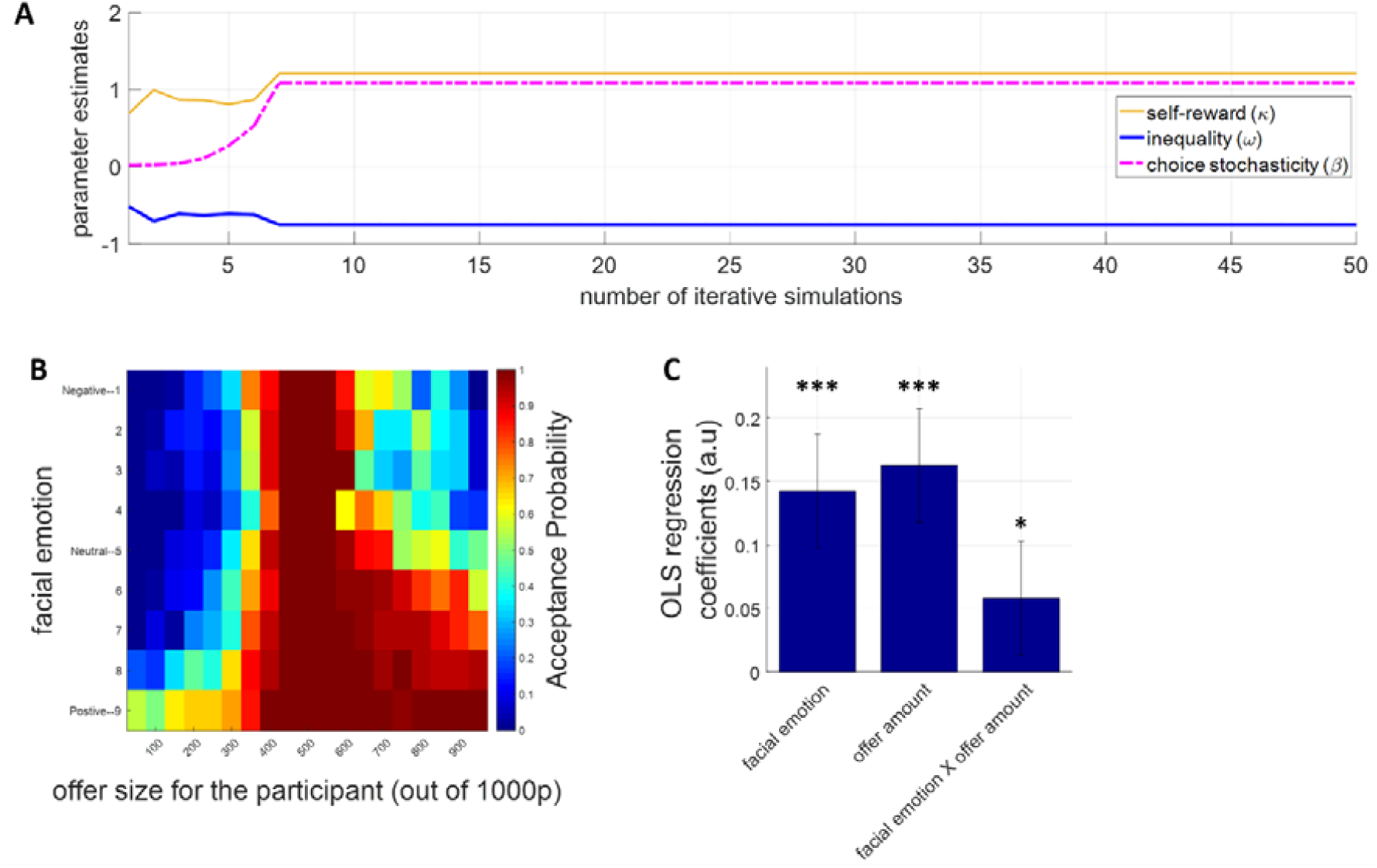
Model recovery analyses. **(A)** Iterative simulations of the best-fitting model based on the population average of the model parameters on a task environment experienced by a representative subject demonstrate that parameter estimates quickly stabilise. In these simulations choices were generated stochastically by a sigmoid function and the parameters in the current simulation (i+1) were re-estimated based on the generated choices from the previous simulation (i). **(B-C)** Reanalysis of the raw data originally reported in Figure 3 by replacing participant choices with the choices generated by the best fitting dynamic inequality aversion model, is able to reproduce the essence of human behaviour and recapture significant main effects of facial emotions, offer amounts and their interaction (*P<0.05, ***P<.001).

**Supplementary Figure 6.**
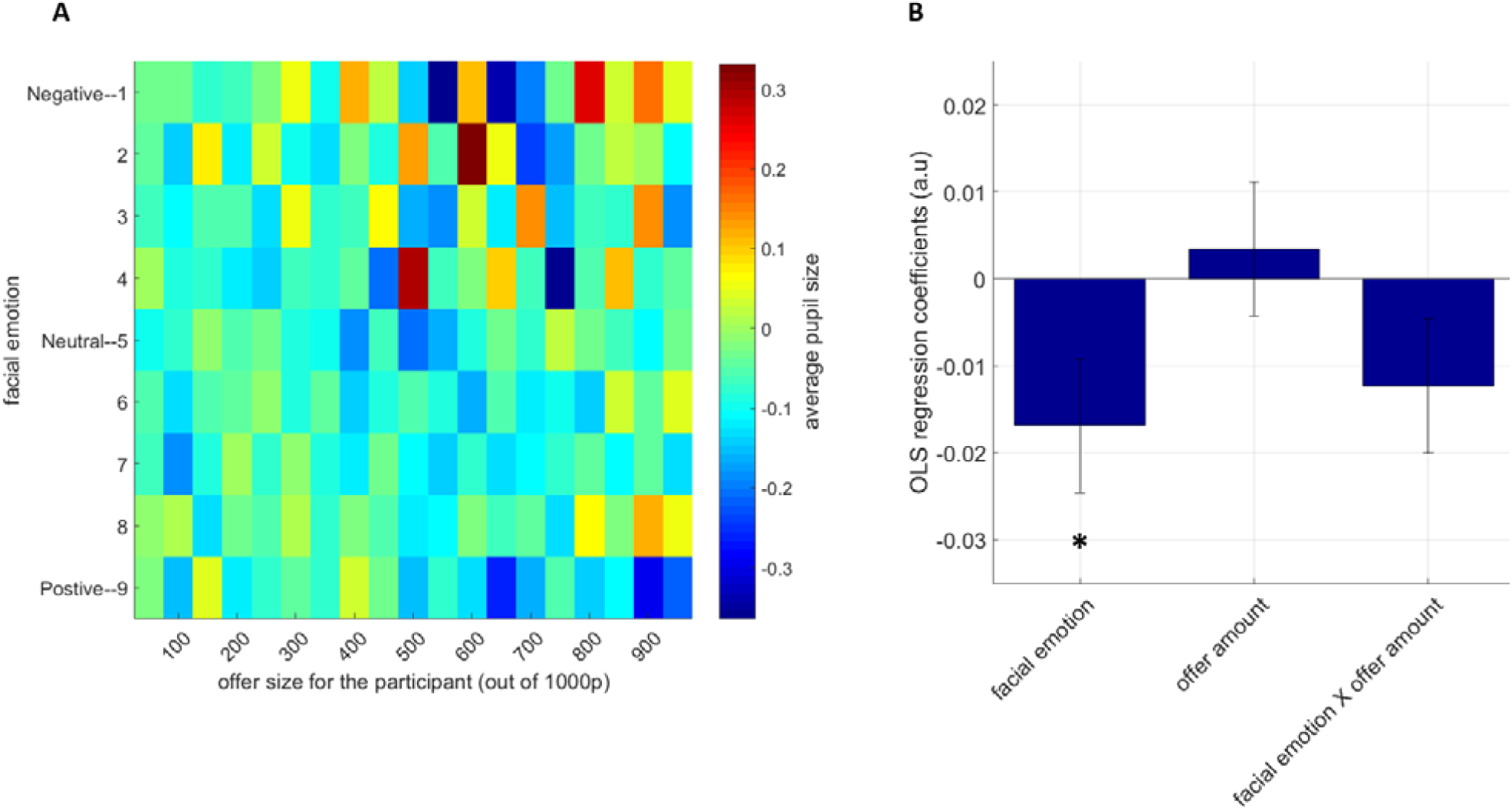
Average pupil size in the Ultimatum Game. **(A).** Participants’ average pupil size across all possible combinations of proposer’s facial emotion (y-axis) and offer amounts (x-axis) represented as a heat map. Colour bar shows the average pupil size after presentation of the offer. **(B)** An OLS regression analysis conducted on the pupil size suggested only a significant main effect of proposer’s facial emotion (*p<.05).

**Supplementary Figure 7.**
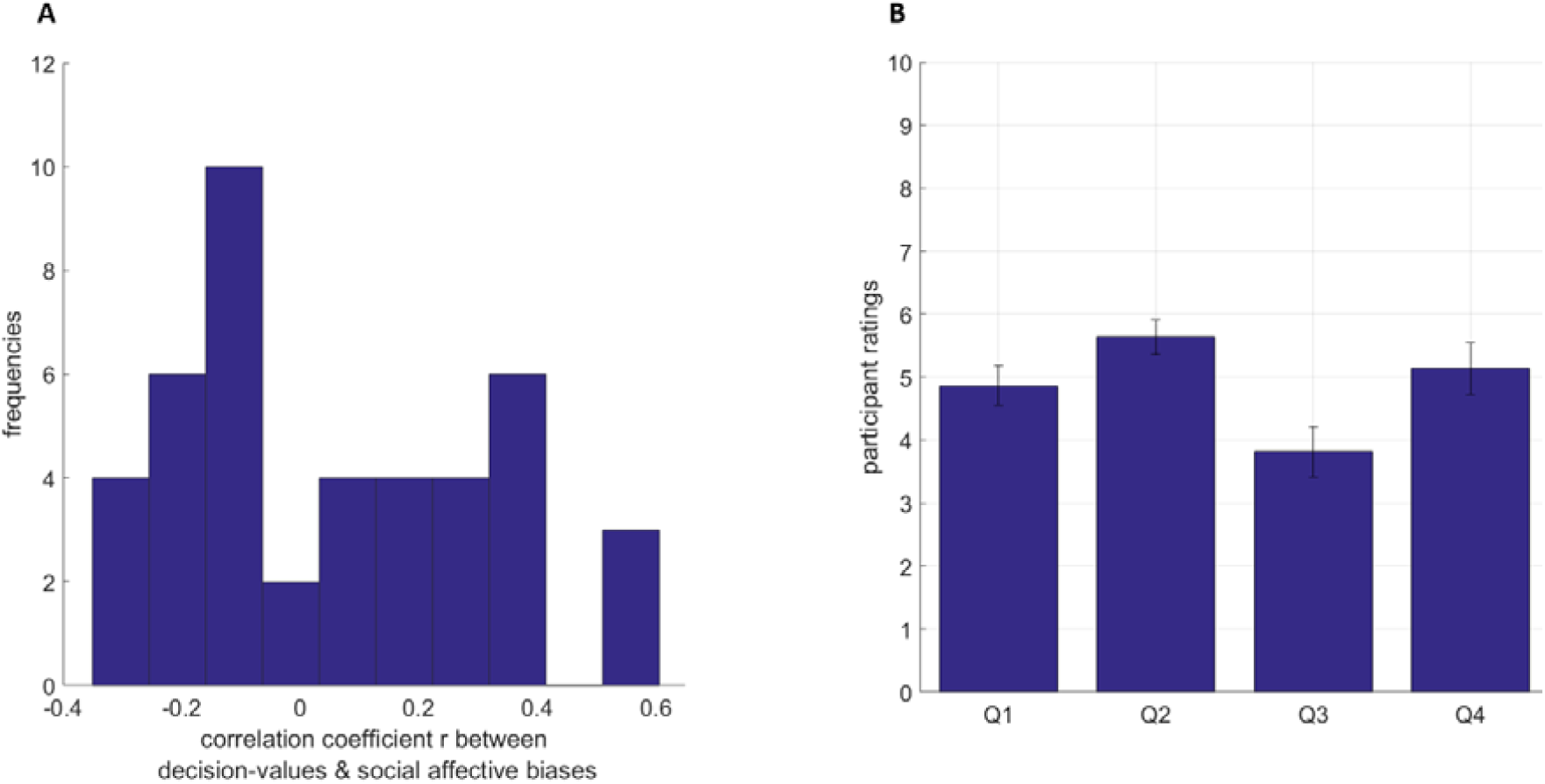
Relationship between social affective biases and decision-values. **(A)** The distribution of correlation coefficients r between social affective biases and decision-values entered in to the pupillary model. Frequencies on the y-axis indicate the number of participants. **(B)** Participants rated the proposer on 4 questions from 0 to 10. Q1. How much do you think this person cares about rewards to others? (0: does not care at all, 10: cares very much) Q2. How much would you like this person if you spent 1 hour with him/her in real-life? (0: strong dislike, 10: very likeable) Q3. How many people do you know in real life who resembles this person (not physical appearance)? (number 0-10) Q4. How socially close do you feel towards those people that you know? (0: not close at all, 10: very close).

**Supplementary Figure 8.**
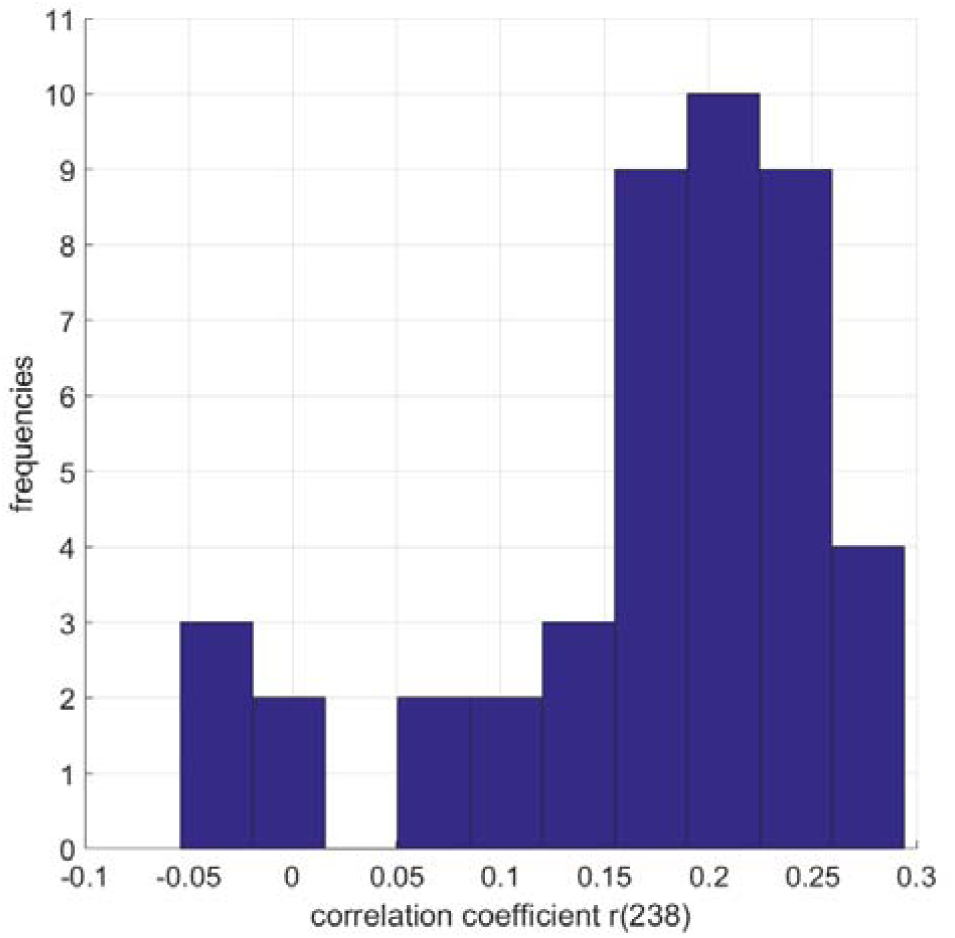
Relationship between surprise associated with offers and the actual offer amount. Distribution of correlation coefficients between the surprise regressor generated by the Bayesian filter uniquely for each participant and the vector of offer amounts observed by the participant throughout the task. Frequencies on the y-axis indicate the number of participants.

